# Human USP18 is regulated by miRNAs *via* the 3’UTR, a sequence duplicated in lincRNA genes residing in chr22q11.21

**DOI:** 10.1101/2020.10.07.328385

**Authors:** Erminia Rubino, Melania Cruciani, Nicolas Tchitchek, Anna Le Tortorec, Antoine D. Rolland, Önay Veli, Leslie Vallet, Giulia Gaggi, Frédérique Michel, Nathalie Dejucq-Rainsford, Sandra Pellegrini

**Affiliations:** Unit of Cytokine Signaling, Institut Pasteur, INSERM U1221, Paris, France; École Doctorale Physiologie, Physiopathologie et Thérapeutique, ED394, Sorbonne Université, Paris, France; i3 research unit, Hôpital Pitié-Salpêtrière-Sorbonne Université, Paris, France; UMR_S1085, Irset (Institut de recherche en santé, environnement et travail), EHESP, Inserm, Univ Rennes, F- 35000 Rennes, France

**Author notes:** Corresponding author : Sandra Pellegrini, Institut Pasteur, 25 rue du Docteur Roux, 75724 Paris cedex 15, France.

**Keywords:** type I IFN, USP18, 3’UTR, lincRNA, chr22q11.21, testis

## Abstract

Ubiquitin-specific peptidase 18 (USP18) acts as gatekeeper of type I interferon (IFN) responses by binding to the IFN receptor subunit IFNAR2 and preventing activation of the downstream JAK/STAT pathway. In any given cell type, the level of USP18 is a key determinant of the output of interferon-stimulated transcripts. How the baseline level of USP18 is finely tuned in different cell types remains ill defined. Here we explored post-transcriptional regulation of USP18 by microRNAs (miRNAs) and identified four miRNAs (*miR-24-3p, miR-191-5p, miR-423-5p* and *miR-532-3p*) that efficiently target *USP18* through binding to the 3’UTR. Among these, three miRNAs are particularly enriched in circulating monocytes which exhibit low baseline *USP18*. Intriguingly, the *USP18* 3’UTR sequence is duplicated in human and chimpanzee genomes. In human, we found several copies of the 3’UTR that are embedded in long intergenic non-coding (linc) RNA genes residing in chr22q11.21 and exhibiting a tissue-specific expression pattern. Interestingly, one of these lincRNAs (here named *linc-UR-B1*) is uniquely and highly expressed in testis. RNA-seq data analyses from testicular cell subsets revealed a positive correlation between *linc-UR-B1* and *USP18* expression in spermatocytes and spermatids. Overall, our findings uncover a set of miRNAs and lincRNAs, which may be part of a network evolved to fine-tune baseline USP18, particularly in cell types where IFN responsiveness needs to be tightly controlled.

**SIGNIFICANT STATEMENT:** USP18 is a non-redundant negative feedback regulator of type I IFN signaling and a key determinant of cell responsiveness to IFN. How baseline USP18 is set in different human cell types is ill defined. We identified three microRNAs that restrain USP18 level notably in primary monocytes through binding the 3’UTR. We found several copies of the USP18 3’UTR embedded in long intergenic non-coding (linc) RNAs which reside in a complex region of human chromosome 22. These lincRNAs are expressed in a tissue-specific manner. We describe one lincRNA expressed only in testis, and most notably in germ cells. Correlative analyses suggest that microRNAs and lincRNAs may form a network controlling baseline USP18 and IFN responsiveness.

## INTRODUCTION

Ubiquitin-specific peptidase 18 (USP18) is an interferon-stimulated gene (ISG) exerting a specific and non-redundant role in the negative regulation of type I IFN (here IFN) responses (1). USP18 is recruited to the IFNAR2 receptor subunit, interferes with JAK1/TYK2 activation, and attenuates STAT-mediated ISGs induction. USP18 is also an isopeptidase (de-ISGylase) that non-redundantly cleaves the ubiquitin-like ISG15 from protein conjugates (2-4). Deficiency in *USP18* causes a severe type I interferonopathy resulting in perinatal death with serious brain malformations due to spontaneous microglia activation, which most likely results from unrestrained response to constitutive IFNβ (5). Indeed, high baseline USP18 maintains microglia quiescence and prevents sub-threshold activation (6, 7). In mouse models of VSV and LCMV infection, it was shown that the ability of CD169+ spleen macrophages and dendritic cells to present viral antigens and elicit an innate and adaptive immunity relies on the expression of *Usp18* (8, 9). Altogether, these findings point to USP18 as a key determinant of cell responsiveness to IFN, including in the context of constitutive low IFNβ levels. We and others have shown that in humans, but not in mice, free ISG15 sustains the level of USP18, by preventing its proteasomal degradation by the S-phase kinase-associated protein 2 (SKP2) (10-12). Yet, little is known on how baseline USP18 levels are set in different cell lineages.

MicroRNAs (miRNAs) act as fine-tuners of gene expression and protein output. Each gene target can be regulated by numerous miRNAs, which typically bind to specific sites in 3’ untranslated regions (3’UTRs) of mRNAs, influencing mRNA stability and translation. Hence, the amount of target varies from cell to cell depending on the identity and the expression level of each miRNA (13). MiRNAs operate in different physiological and pathological processes (14) and also participate to the complex network that regulates immune cell development, function and response to stimuli (15). As for many immune pathways, miRNAs have been shown to regulate the IFN system both at the level of production and response. Hence, some of the key players of the type I IFN signaling pathway are under the control of miRNAs (16), but no miRNA targeting USP18 has been described so far.

Here, we investigated whether miRNAs contribute to tune down baseline USP18 in a cell-context specific manner. We identified four miRNAs that negatively regulate USP18 through binding the 3’UTR. We also report on the presence of several copies of the *USP18* 3’UTR in the human genome. These duplications reside on chr22q11.21 and are embedded in long intergenic non-coding RNA (lincRNA) genes, whose expression is tissue-specific. The remarkable cell type-specific expression of one such lincRNA in testicular germ cells and its positive correlation with *USP18* raises the possibility that the *USP18* 3’UTR not only acts as a *cis*-regulatory sequence targeted by miRNAs, but also acts in *trans* when embedded in lincRNA molecules expressed in particular cell types.

## RESULTS

### The 3’UTR of human USP18 is targeted by at least four miRNAs

Using the bioinformatic prediction programs miRWalk and RNAhybrid, we identified 27 miRNAs predicted to target with high score the 580 nucleotide (nt)-long 3’UTR of the human *USP18* mRNA (**Fig. 1A**). *USP18* is basally detected in most human tissues, albeit at variable levels (**Fig. S1**). We therefore analyzed the expression of the 27 miRNA candidates in the 90 cell types of the FANTOM5 dataset (17). Fourteen miRNAs were found to be expressed at > 50 CPM (counts per million mapped reads) in at least one cell type (**Fig. 1A**). We next tested the functional impact of these 14 miRNAs on USP18. For this, we expressed in HeLa S3 cells each miRNA as mimic and measured the level of *USP18* mRNA by qPCR. As shown in **Fig. 1B**, seven miRNAs led to a reduction of endogenous *USP18*, four of them exhibiting a stronger effect. Consistently, this also resulted in lower USP18 protein levels, at least for those four miRNAs with stronger effects at the RNA level (**Fig. 1C**).

**Figure 1.**
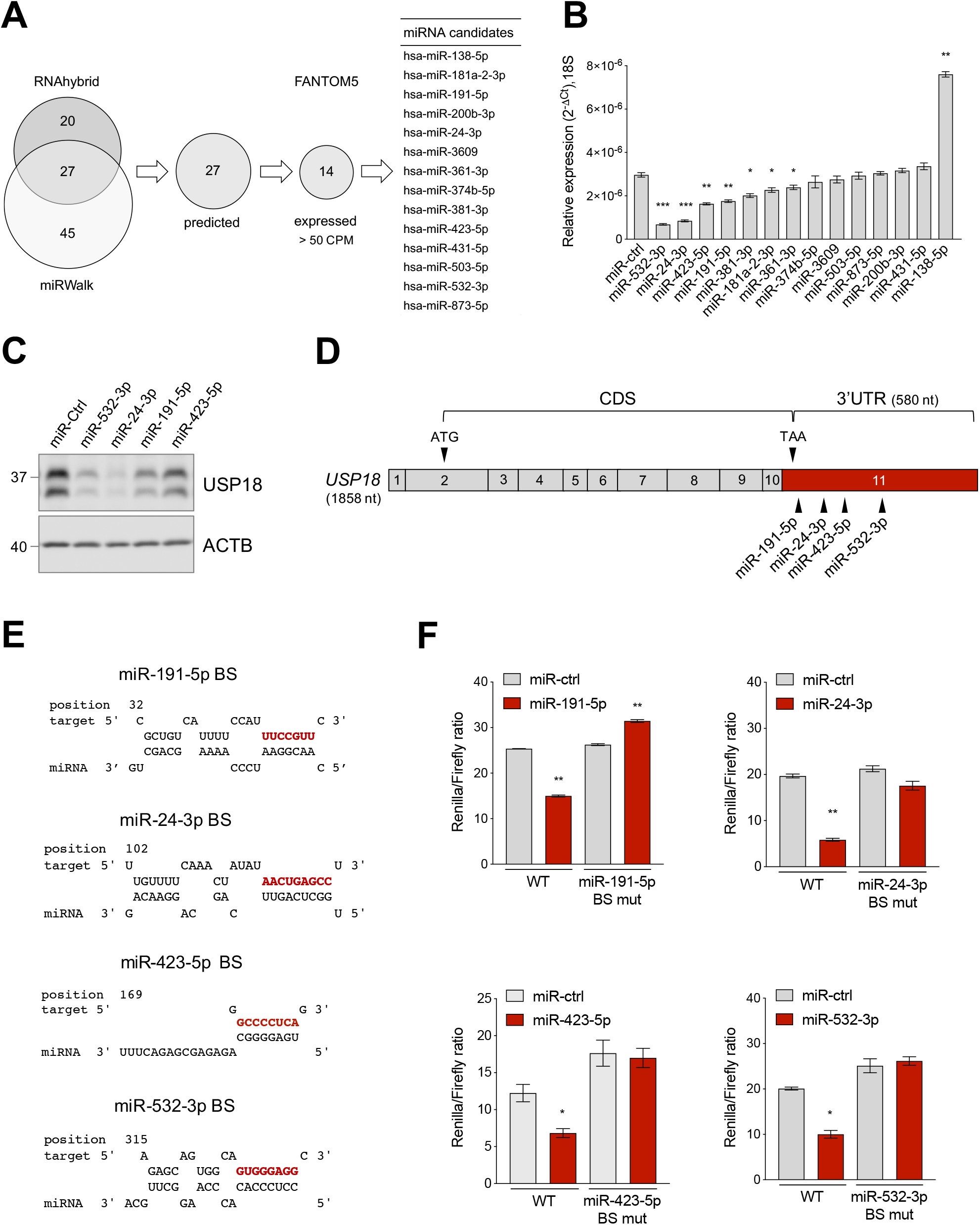
Identification of USP18-targeting miRNAs. **A)** Prediction of *USP18*-targeting miRNAs using RNA hybrid algorithm and mirWalk software (TargetScan, Pita, miRDB, RNA22). The common high-scoring miRNAs (27) were further selected for expression (counts per million mapped reads, CPM > 50) in 90 human cell types (FANTOM5 dataset). The selected candidates (14) are listed. **B)** The 14 selected miRNAs and a miR-control (miR-ctrl) were expressed as mimics in HeLa S3 cells and *USP18* mRNA was measured by qPCR (n = 4). Results (± SEM) shown as expression (2^-ΔCt^) relative to *18S*, used as housekeeping gene. Raw-matched one-way ANOVA with Geisser-Greenhouse correction was performed (Dunnett’s multiple comparison - compared to miR-ctrl), adjusted p-values shown as stars * < 0.05, ** < 0.01,*** < 0.001, **** < 0.0001. **C)** The miR-ctrl and the indicated miRNAs were expressed as mimics in HeLa S3 cells and 48 hr later endogenous USP18 was measured by western blot. Actin B (ACTB) as loading control. The data are representative of three experiments. **D)** Schematic of the *USP18* mRNA. Numbered boxes correspond to the exons. CDS, coding sequences. Arrowheads, the start (ATG) and stop (TAA) codons and the binding sites for the four indicated miRNAs. Note that exon 11 (in red) contains a short coding sequence (43 nt) and the stop codon. **E)** Binding sites (BS) of *miR-191-5p, miR-24-3p, miR-423-5p* and *miR-532-3p* in the *USP18* 3’UTR. Seed-matched sequences are in red. Pairing of miRNA-*USP18* sequences was obtained with the RNAhybrid algorithm. **F)** HeLa S3 cells were transfected with the indicated miRNAs and 24 hr later with the psiCHECK2-*USP18* 3’UTR reporter plasmid, WT or mutated in the miRNA binding site (BS mut). Each mutant plasmid contains a 5 nt mutation (nt 2 to 6 of the seed-matched sequence shown in E). Luciferase activity was measured 24 hr after plasmid transfection (n = 4) and expressed as Renilla/Firefly ratio (± SEM).

In order to validate the binding of each miRNA to the predicted site on *USP18* 3’UTR (**Fig. 1D, E**), this latter sequence was subcloned downstream of the *Renilla* luciferase gene in the psiCHECK2 vector and the impact of the four miRNAs on luciferase was measured in transfected HeLa S3 cells. As shown in **Fig. 1F**, *miR-191-5p, miR-24-3p, miR-423-5p* and *miR-532-3p*, expressed as mimics, significantly decreased luciferase activity with respect to the control mimic. Moreover, the effect of each mimic was abrogated when a 5 nt mutation was introduced in the seed-matched sequence of the predicted binding site on the 3’UTR (**Fig. 1F**). Altogether these data demonstrate that the *USP18* 3’UTR sequence can be directly targeted by *miR-191-5p, miR-24-3p, miR-423-5p* and *miR-532-3p*, with an effect on USP18 abundance.

To determine in which cell type(s) this post-transcriptional control may operate, we performed a principal component analysis (PCA) based on the expression of the seven active miRNAs identified above in the 90 cell types (**Fig. 2A**). The analysis revealed a different distribution of immune *vs* non-immune cells, with the first three PC vectors explaining 75% of the total variance. While the distribution of non-immune cells was homogenous, the distribution of immune cells was more scattered and mostly driven by *miR-191-5p, miR-24-3p, miR-532-3p* and *miR-361-5p* (**Fig. 2A**). These four miRNAs were expressed variably across the 90 cell types, but were enriched in immune cells (**Fig. S2A**). We therefore performed a second PCA based on expression of these four miRNAs exclusively in circulating peripheral blood mononuclear cells (PBMCs) (CD14^+^ monocytes, CD8^+^ T cells, CD4^+^ T cells, NK cells, CD19^+^ B cells). These miRNAs appeared to contribute to the distribution of the cell populations analyzed (**Fig. 2B)**. In particular, *miR-191-5p, miR-24-3p* and *miR-532-3p* were found to be enriched in monocytes, which clustered apart from all lymphoid cells **(Fig. S2B**). To reinforce these data, we surveyed an independent dataset (Cohort Roche, GSE28492) (18), which confirmed a higher level of the three miRNAs in monocytes with respect to NK, T and B cells (**Fig. 2C**). Of note, *miR-191-5p* and *miR-24-3p* were more abundant than *miR-532-3p* in monocytes.

**Figure 2.**
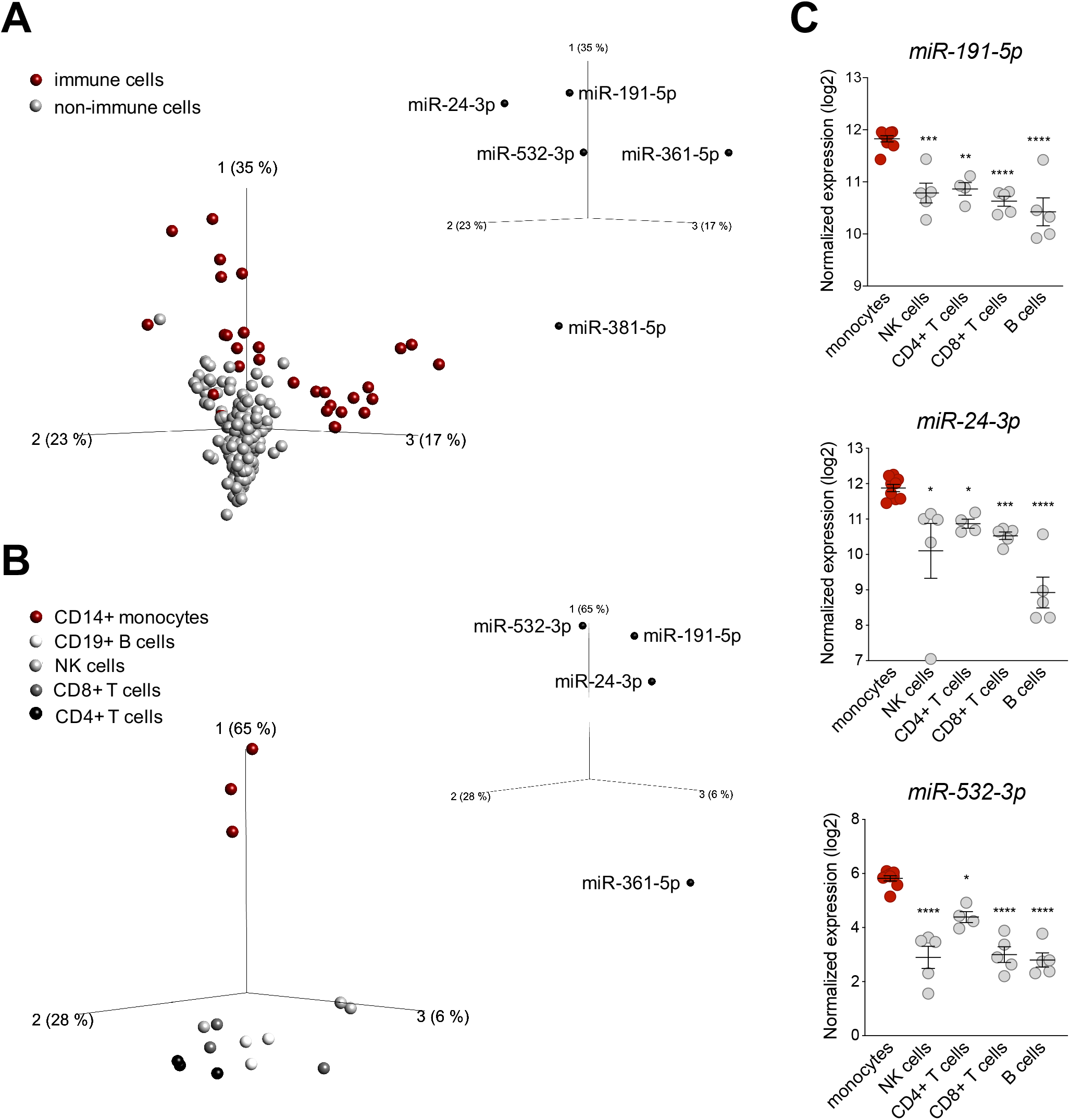
*miR-191-5p, miR-24-3p* and *miR-532-3p* are enriched in monocytes. **A)** Principal component analysis (PCA) performed on 90 human cell types (FANTOM5 project dataset, 3 donors per cell population). The distribution is based on expression of the seven miRNAs significantly targeting *USP18* (see Fig. 1B). Red and gray circles: immune and non-immune cells, respectively. The PCA plot shown captures 75% of the total variance within the selected data set (PCA1 35%; PCA2 23%; PCA3, 17%). q-value (ANOVA FDR adjusted p-value) < 0.05. The five *USP18*-targeting miRNA candidates significantly contributing to the data distribution are shown on the three PC axes of the plot. **B)** PCA performed on circulating immune cell populations (FANTOM5 project data set). Analysis was restricted to the four *USP18*-targeting miRNAs enriched in immune cells. Color legend on the left. The PCA plot captures 97% of the total variance within the selected data set (PCA1 65%; PCA2 28%; PCA3 6%). q-value (ANOVA FDR adjusted p-value) < 0.05. The four miRNAs significantly contributing to the data distribution are shown on the three principal component axes of the PCA plot. **C)** Normalized expression (log2, ± SEM) of *miR-532-3p, miR-191-5p* and *miR-24-3p* in circulating immune cell subsets. Data retrieved from (18), Cohort Roche, GSE28492. Ordinary one-way ANOVA (Dunnett’s multiple comparison - compared to monocytes) was performed, adjusted p-values shown as stars * < 0.05, ** < 0.01, ***< 0.001.

The enrichment of *miR-191-5p, miR-24-3p* and *miR-532-3p* in monocytes prompted us to search for a correlation with *USP18*. For this, CD14+ monocytes and monocyte-depleted PBMCs (here PBL for peripheral blood lymphocytes) were freshly isolated from blood of four donors and levels of the three miRNAs and of *USP18* were measured by qRT-PCR. The three miRNAs were more expressed in monocytes than PBL (**Fig. 3A**), while *USP18* exhibited an opposite profile (**Fig. 3B**). In addition, a significant negative correlation between each miRNA and *USP18* was observed (**Fig. 3C**). Interestingly, the opposite was observed for four other ISGs, *i.e. OAS1, IRF7, IFIT1* and *STAT2*, which were found to be more abundant in monocytes than PBL, suggesting a specific profile for USP18 as compared to other ISGs (**Fig. 3D**).

**Figure 3.**
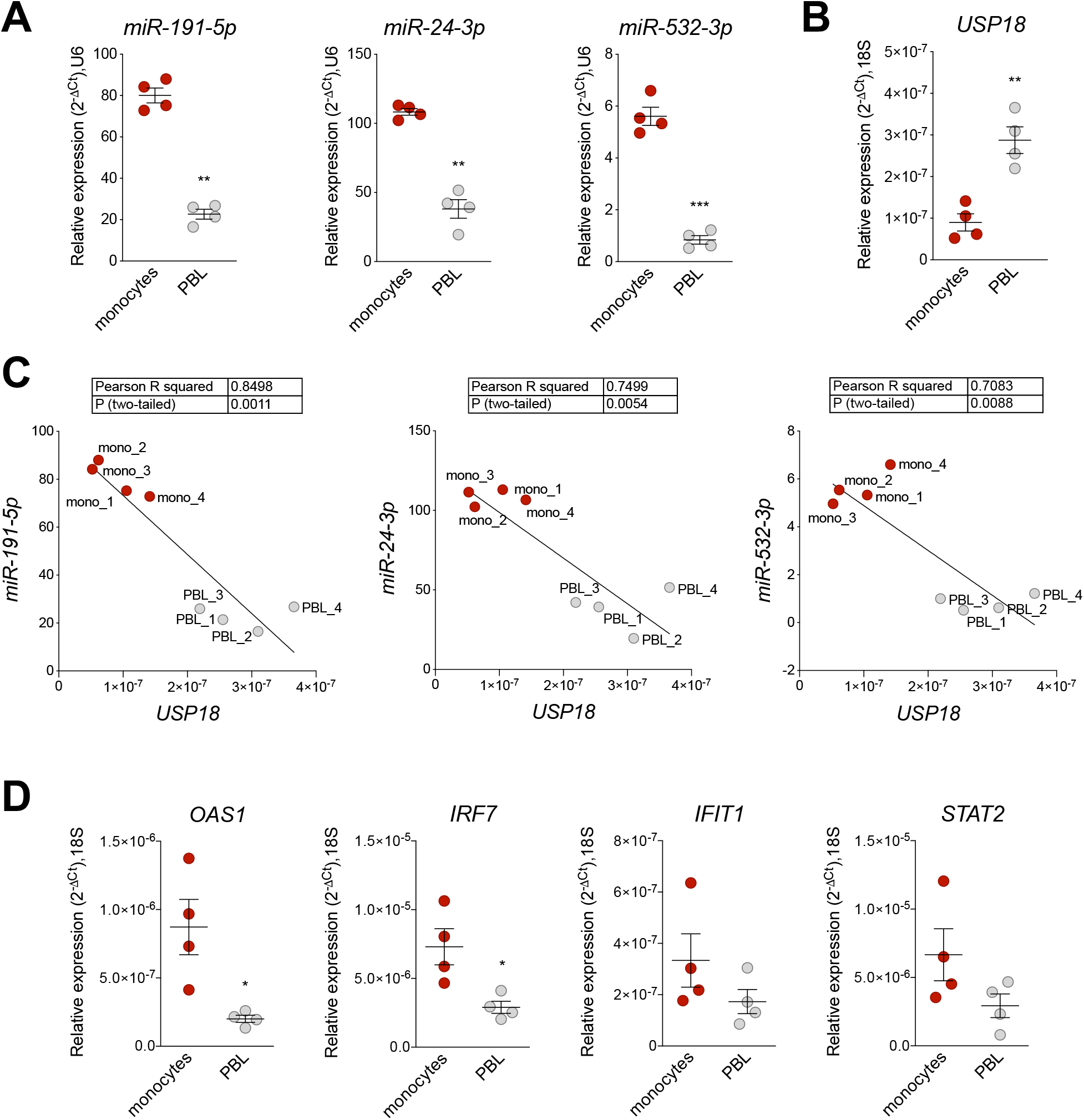
*miR-191-5p, miR-24-3p* and *miR-532-3p* negatively correlate with *USP18* in monocytes *vs* PBL. **A)** and **B)** Expression of *miR-191-5p, miR-24-3p* and *miR-532-3p* (A) and of *USP18* (B) in *ex vivo* monocytes and monocyte-depleted PBMCs (here PBL) (four donors). Results (± SEM) shown as expression (2^-ΔCt^) relative to *U6* or *18S*, used as housekeeping genes. Paired t test p-values shown as stars ** < 0.01, *** < 0.001. **C)** Negative correlation between *USP18* and *miR-191-5p, miR-24-3p and miR-532-3p* in monocytes and PBL. Expression levels used to calculate the correlation are shown in A) and B). Pearson correlation coefficients and p-values (two-tailed) are shown above. **D)** Expression of *OAS1, IRF7, IFIT1* and *STAT2* in monocytes and PBL (4 donors). Results (± SEM) shown as expression (2^-ΔCt^) relative to *18S*. Paired t test p-values are shown, when significant, as stars * < 0.05.

Altogether, these results suggest that *miR-191-5p, miR-24-3p*, and *miR-532-3p* contribute to restraining baseline *USP18* in human monocytes. Although not enriched in monocytes as compared to PBL, *miR-423-5p* may also contribute as it was relatively highly expressed among the studied miRNAs (**Fig. S3**).

### Copies of USP18 3’UTR embedded in lincRNA genes

The results described above suggest that the level of human USP18 can be post-transcriptionally fine-tuned by four miRNAs binding its 3’UTR. While these miRNAs and their genes are conserved down to the mouse (**Fig. S4A**), their binding sites on the 3’UTR are not. In fact, seed-matched sequences for only two of them (*miR-24-3p* and *miR-532-3p*) are conserved in the *USP18* 3’UTR of non-human primates (**Fig. S4B, C**). In searching for sequence conservation, we found several copies of the *USP18* 3’UTR in the human and chimpanzee genomes (**Table 1**), raising the possibility that transcripts other than the *bona fide* USP18 protein-coding mRNA contain the *USP18* 3’UTR and bind the same miRNAs.

**Table 1.**
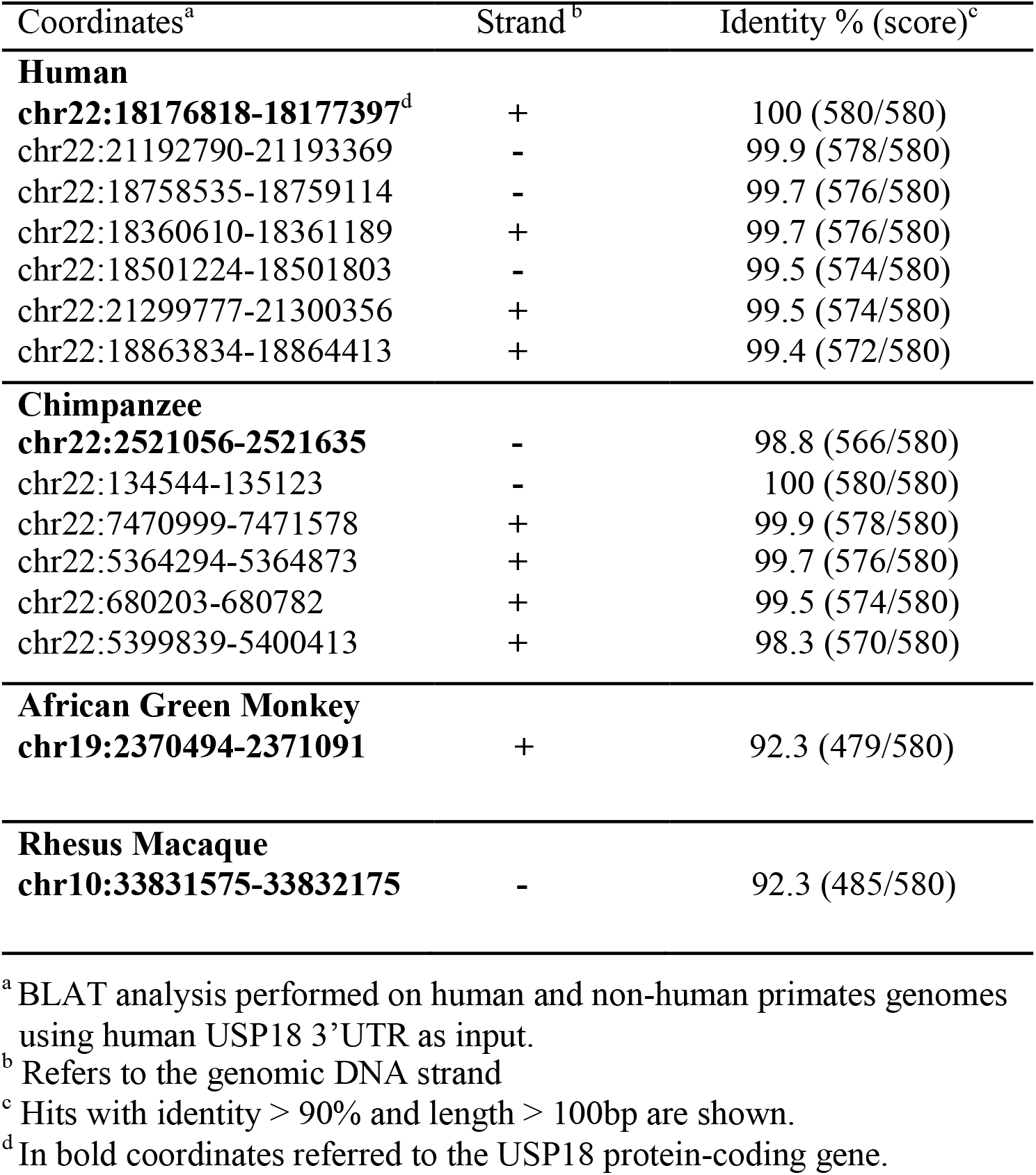
*USP18* 3’UTR copies in human and non-human primates.

The *USP18* gene and all copies of the 3’UTR sequence map on chr22q11.21, a highly complex region of the human genome that contains four segmental duplications or low-copy repeats (LCR22-A to D) (**Fig. 4A**) (19). The high sequence identity and copy number variation in the LCR22s have indeed hampered accurate sequencing and gene annotation in this region and, even in the most recent version of the human reference genome, LCR22s are rich in gaps and assembly errors (20). By performing a BLAT analysis of *USP18* gene sequences on the last hg38 assembly, we identified six copies of the 3’UTR in positive or negative strand orientation (**Fig. 4A** and **Table 1**). Four copies - named A1 to A4 here - contain most of intron 10 and the entire exon 11 (*i.e*. the *USP18* 3’UTR) of the *USP18* gene. The other two copies, named B1 and B2, contain a small segment of intron 10 and the entire exon 11. All six copies reside in LCR-A and LCR-D (**Fig. 4A, B**). Importantly, with the exception of B2, the other copies overlap each an annotated long intergenic non-coding RNA (lincRNA) gene (see list in **Table 2**). The lincRNA genes spanning copies A1, A3 and A4 encode three annotated XR_transcripts of nearly identical sequence, which we named *linc-UR-A1, linc-UR-A3* and *linc-UR-A4* respectively, where UR stands for USP18-Related (**Fig. 4C**). The annotated transcript spanning A2 (*linc-UR-A2*) contains only few nucleotides of the 3’UTR and was not further studied. The lincRNA gene spanning B1 (named *linc-UR-B1*) encodes two transcript isoforms (annotated as *TCONS_00029753* and *TCONS_00029754*) differing of 4 nt at the exon-exon junction (**Fig. 4D**) (**Fig. S5A**). As annotated, these transcripts contain two exons: exon 1 differing between *linc-UR-A1/3/4* and *linc-UR-B1* and exon 2 corresponding to *USP18* exon 11 (**Fig. 4C, D**).

**Table 2.**
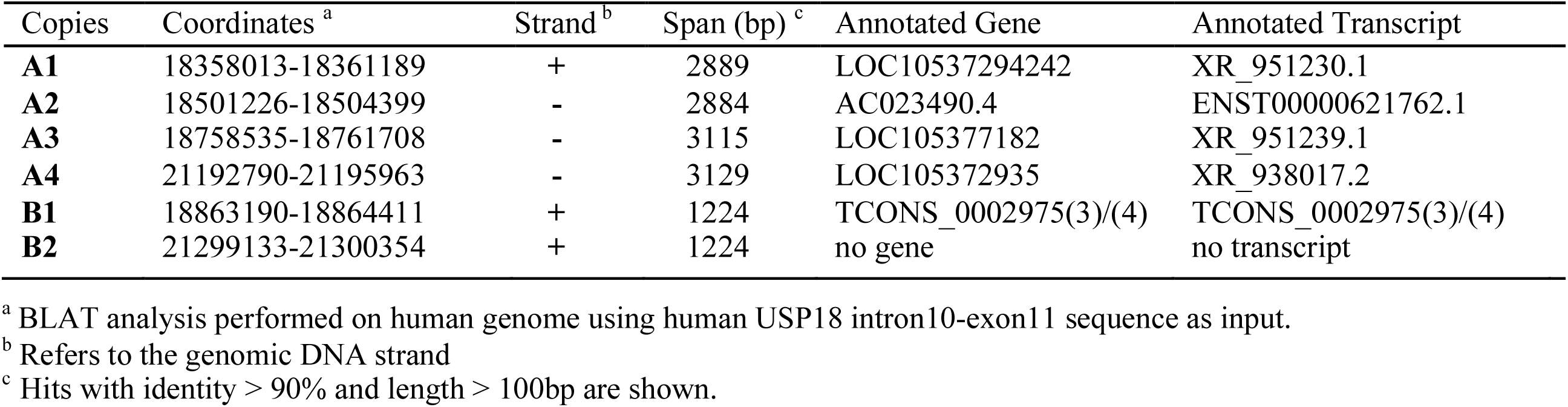
Copies of USP18 3’UTR embedded in lincRNA genes.

**Figure 4.**
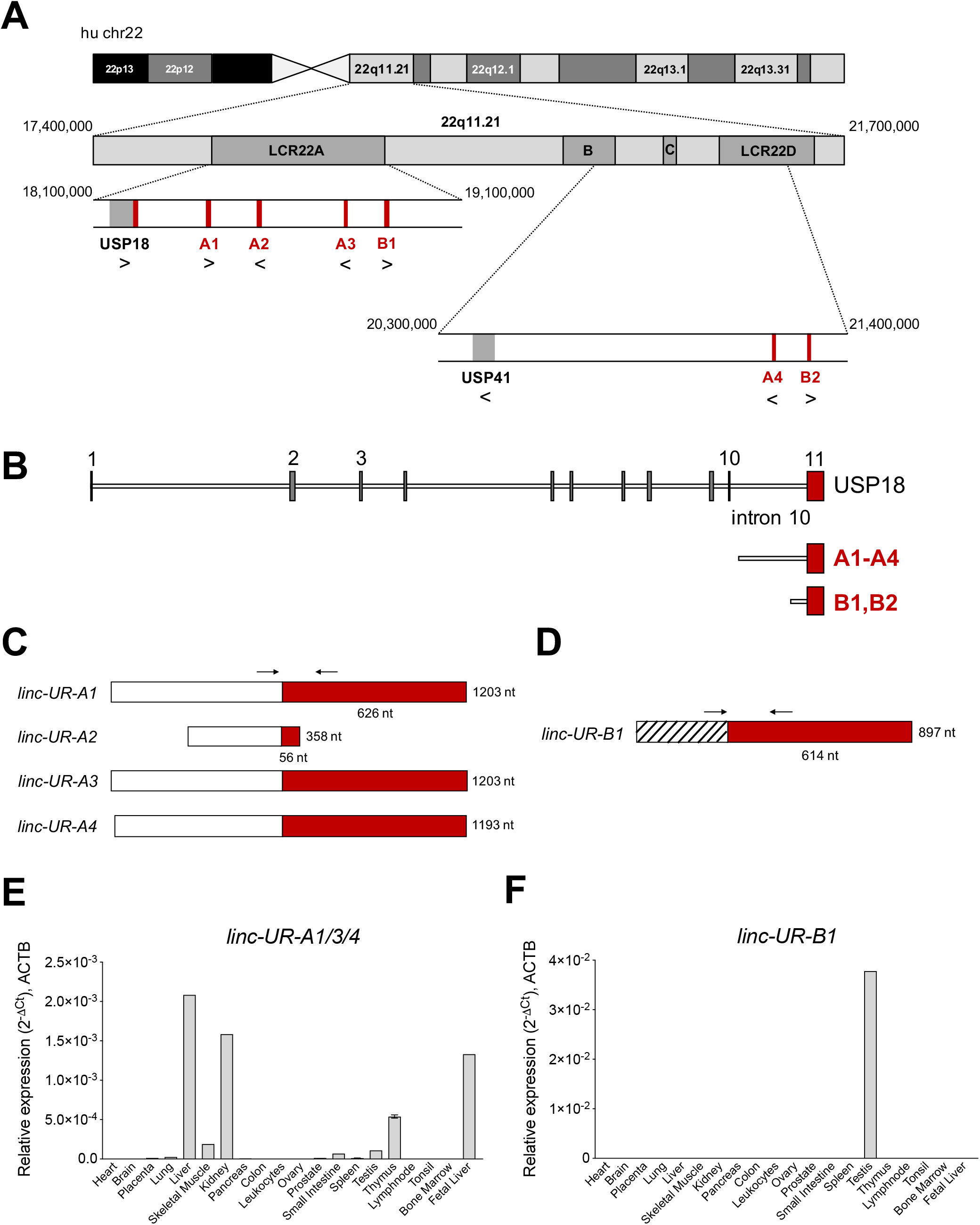
Several *USP18* exon 11 copies are embedded in expressed lincRNA genes. **A)** Schematic of human chr22 (about 50 Mb) and below the chr22q11.21 region with four LCR22s. The *bona fide USP18* gene resides at the boundary of LCR22A. The pseudogene *USP41*, located in LCR22B, contains most *USP18* gene sequences, *i.e*. from exon 3 to exon 10, but lacks the 5’ and 3’ UTRs. The six copies of *USP18* exon 11 are indicated in red. Genes and *USP18* copies are shown with their genomic orientation (> for + strand, < for - strand). **B)** Top, intron-exon organization of the human *USP18* gene. Exons, grey boxes; introns, lines. Exon 11 is highlighted in red. Below are aligned the duplications found. Copies A1 to A4 contain most of *USP18* intron 10 and the entire exon 11. Copies B1 and B2 contain a smaller sequence of intron 10 and exon 11. **C)** Map of *linc-UR-A* transcripts (*A1* to *A4*) containing *USP18* exon 11 in red. The annotated exon upstream of exon 11 is represented as white box. Arrows indicate the primers (A_FW1 and A_RV1) designed across the junction and used to measure expression of *linc-UR-A* in (E). **D)** Map of *linc-UR-B1*. The annotated exon upstream of exon 11 is represented as striped box. Arrows indicate the primers (B_FW1 and B_RV1) designed across the junction and used to measure expression of *linc-UR-B1* in (G). **E)** Expression of *linc-UR-A1/3/4* in 20 human tissues (pool of donors for each tissue; Human Immune System MTC™ Panel, Human MTC™ Panel I, Human MTC™ Panel II). Results shown as expression (2^-ΔCt^) relative to *ACTB*. SEM is shown for tissues present in two of the above panels. **F)** Expression of *linc-UR-B1* in the same samples used in (E).

Next we measured the expression of these transcripts by qPCR using a commercial panel of cDNAs from polyA+ RNAs of 20 human tissues. Importantly, we designed specific primers across the unique exon-exon junction of the *linc-UR-A* transcripts (A1, A3 and A4) and across the unique exon-exon junction of *linc-UR-B1* (**Fig. 4C, D**) (primers in **Table S2**). *Linc-UR-A* transcripts were lowly expressed in most tissues and showed the highest expression in fetal and adult liver, kidney, and thymus (**Fig. 4E**). On the other hand, *linc-UR-B1* was uniquely and abundantly expressed in testis (**Fig. 4F**). To confirm the testis-restricted expression of *linc-UR-B1*, we interrogated RNA-seq datasets covering a panel of 32 tissues (E-MTAB-2836). To distinguish *linc-UR-B1* from all other *USP18*-containing transcripts, uniquely mapped reads were selected and aligned again to the human genome (hg38). This analysis revealed that *linc-UR-B1* is expressed only in testis as a single isoform corresponding to the *TCONS_00029754* annotation (**Fig. S5B**). This was further confirmed by sequencing the only product amplified by qPCR on testis cDNA (**Fig. S5C**). High inter-individual variability in the level of *linc-UR-B1* was noticed in the RNA-seq dataset (eight donors) (**Fig. S5B**). Our qPCR analyses on testis fragments from six donors displaying normal spermatogenesis and one donor with impaired spermatogenesis (as observed by transillumination of seminiferous tubules) confirmed this variability and showed the lowest *linc-UR-B1* expression in the latter donor (**Fig. S5D**).

As opposed to protein-coding genes, the annotations of lincRNAs are far from being complete (21). To validate the existing annotations, we performed 3’ RACE on RNAs from liver and testis. We confirmed the 3’ end of *linc-UR-A* transcripts as annotated (**Fig. 5A**, right panel) and showed that the 3’ end of *linc-UR-B1* extends further than annotated and actually contains the entire *USP18* 3’UTR (580 nt) (**Fig. 5B**, right panel). Mapping the 5’ end of these transcripts proved to be challenging, since the sequences that are annotated as exon 1 (*i.e*. upstream of the 3’UTR) are themselves duplicated, with several copies residing on chromosome 22 and other chromosomes (**Table S1**). Hence, to enrich for transcripts bearing the *USP18* 3’UTR we performed a reverse transcription (RT) PCR reaction on RNAs from liver and testis using a specific primer (spRT) complementary to the 3’UTR and using forward primers specific to the annotated 5’ ends (**Fig. 5A, B**, left panels). These analyses confirmed the annotated sequences for *linc-UR-A* and *linc-UR-B1*. Moreover, we could not amplify *linc-UR-A* further using primers designed in the upstream genomic region, suggesting that this RNA does not extend further.

**Figure 5.**
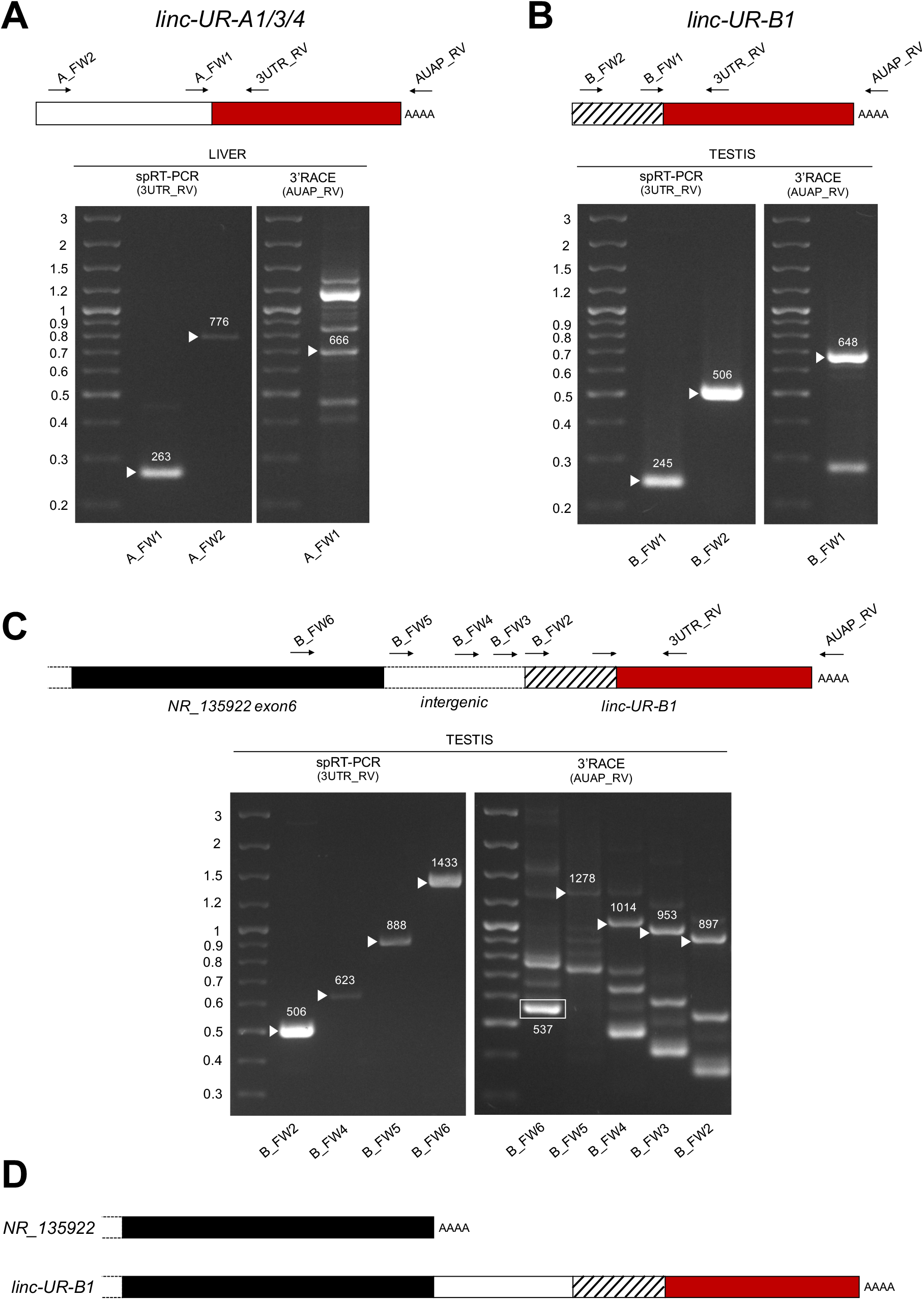
Characterization of *linc-UR-A1/3/4* and *linc-UR-B1* molecules. **A)** Analysis of *linc-UR-A* transcripts in liver. Right gel, 3’RACE. cDNA was obtained by oligo-dT reverse transcription (RT) of polyA+ liver RNA. PCR was performed on this cDNA using the forward (A_FW1) and reverse (AUAP_RV) primers indicated on the schematic above. Several bands were obtained, including the expected 666 bp. Left gel, specific RT-PCR (spRT-PCR). cDNA was obtained from polyA+ liver RNA using a primer designed on the *USP18* 3’UTR (3UTR_GSP1). PCR was then performed using the forward primer (A_FW1 or A_FW2) and a reverse primer designed on the 3’UTR (3UTR_RV) and internal to 3UTR_GSP1. The length in nt is indicated above each amplified product. **B)** Analysis of *linc-UR-B1* transcripts in testis. The strategies described in (A) were used on testis poly A+ RNA. Primers are indicated on the schematic above. The length in nt is indicated above each amplified band. **C)** Study of *linc-UR-B1* transcript extending on the 5’ end. Three forward primers were designed on the annotated intergenic region (B_FW3-4-5). One primer (B_FW6) was designed on the sequence further upstream, corresponding to exon 6 of the annotated *NR_135922* transcript. Right panel, 3’RACE on testis RNA. The 537 nt product obtained using the B_FW6 primer corresponds to the annotated *NR_135922*. Longer products indicated by arrowheads were obtained when using the three primers designed in the intergenic region. Left panel, spRT-PCR was performed on testis RNA using the indicated forward primers and the 3UTR_RV primer. PCR products of the expected lengths were obtained for all primers tested. Panels A), B) and C): the PCR products of the size expected from the annotations were sequenced. The additional 3’RACE PCR products likely result from the amplification of other transcripts containing sequences identical to the exon upstream of *USP18* exon11 (Table S1). **D)** Schematic of the 3’ end of the two *NR_135922* isoforms identified here. Top, the *NR_135922* transcript terminating with exon 6, as in the annotation. Bottom, longer isoform of *NR_135922* terminating with the *USP18* 3’UTR, here called *linc-UR-B1*.

The genomic region upstream of *linc-UR-B1* contains a pseudogene annotated as *NR_135922* and belonging to the large *POM121* family (**Fig. S6A**). Similar to *linc-UR-B1, NR_135922* is expressed only in testis (**Fig. S6B**). To study whether *linc-UR-B1* transcript extends 5’ into *NR_135922* sequences, we designed primers either in the last annotated exon of *NR_135922* or in the intergenic region, and combined them with a reverse primer in the *USP18* 3’UTR. For all primer pairs, the expected products could be amplified from testis RNA (**Fig. 5C** left panel and **5D**), indicating that *NR_135922* and *linc-UR-B1* are part of the same gene. Indeed, 3’RACE on testis RNA yielded the expected *NR_135922* product (537 nt) and a longer isoform (*linc-UR-B1*) that terminates with *USP18* 3’UTR (**Fig. 5C**, right panel and **5D**).

### Linc-UR-B1 is uniquely expressed in male germ cells and positively correlates with USP18

Given the high and specific expression of *linc-UR-B1* in the testis, we sought to identify in which testicular cell type(s) this RNA is expressed. We first performed qPCR analysis on cDNAs from freshly isolated testicular germ cell populations (spermatocytes and spermatids) and from primary somatic peritubular, Leydig and Sertoli cells and detected *linc-UR-B1* only in spermatocytes and spermatids (**Fig. 6A**). To obtain further information on *linc-UR-B1* expression, we surveyed a single-cell RNA-seq dataset of testicular cells from three individuals (22). The expression profile of *linc-UR-B1* was analyzed by tracking the sequence of the unique exon-exon junction between the 3’UTR and the upstream exon. As shown in **Fig. 6B**, *linc-UR-B1* was primarily detected in specific germ cell populations (clusters 4-6). Interestingly, the induction of *linc-UR-B1* expression occured in the transition between meiotic early primary and late primary spermatocytes. This burst was followed by a gradual decrease in post-meiotic round and elongated differentiating spermatids. No or low expression was detected in somatic cell populations (clusters 9-13). To identify more precisely the meiotic phase in which *linc-UR-B1* induction occurs, we analyzed the dataset of early and late primary spermatocytes reclustered for the five different stages of meiotic prophase I. *Linc-UR-B1* was first detectable in the late pachytene stage and was maximal during the diplotene stage (**Fig. 6C**). Importantly, we observed that the expression of *USP18* paralleled that of *linc-UR-B1* in germ cell populations and during prophase I (**Fig. 6D, E**). Since *USP18* is IFN-inducible, its expression may be related to the presence of local IFN. To test this, we analyzed expression of two ISGs, *IFIT1* and *OAS1*, but no or minimal levels of these transcripts were detected in germ cells (**Fig. S7**). In fact, *IFIT1* and *OAS1* were higher in cells other than germ cells, whereas *USP18* was highest in meiotic and post-meiotic germ cells (**Fig. 6D**). The positive correlation between *linc-UR-B1* and *USP18* expression during spermatogenesis (**Fig. 6F**) and in particular in meiotic cells (**Fig. 6G**) suggests that these transcripts - which share the same 3’UTR - may cross-talk. Interestingly, we found that testis, and in particular germ cells, express moderate to high levels of three *USP18*-targeting miRNAs, namely *miR-191-5p, miR-24-3p* and *miR-423-5p* (**Fig. S8**). These data hint at the possibility of a miRNA-dependent cross-talk between *USP18* and *linc-UR-B1* in germ cells, with the latter possibly acting as a sponge for miRNAs targeting USP18.

**Fig. 6.**
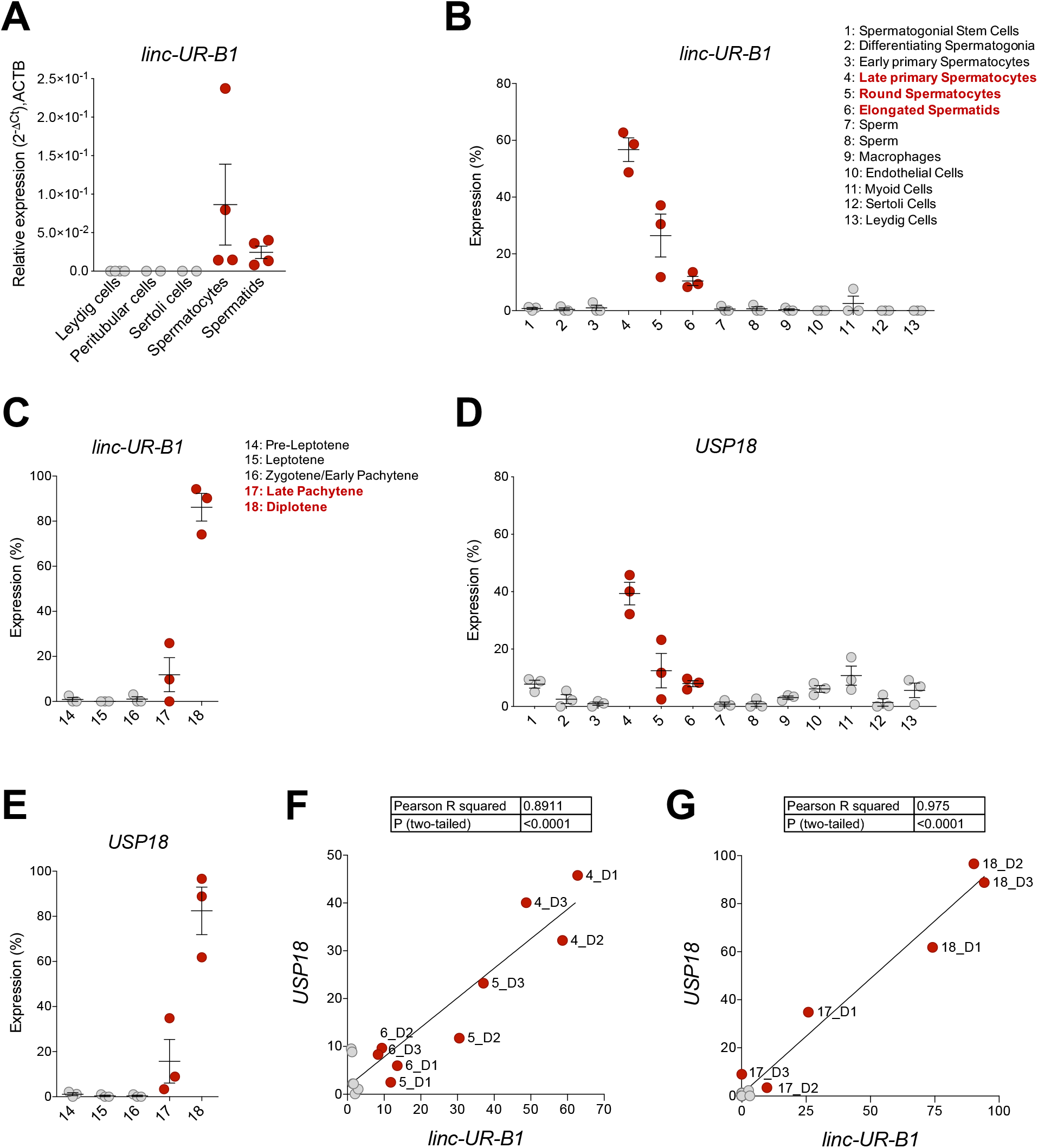
*linc-UR-B1* is expressed in meiotic and post-meiotic germ cells and positively correlates with *USP18*. **A)** Measure of *linc-UR-B1* levels by qPCR in purified testicular cell populations (spermatocytes, spermatids, peritubular cells, Leydig cells) and Sertoli cells. Results (± SEM) shown as expression (2^-ΔCt^) relative to *ACTB*. **B)**and **D)** Analysis of *linc-UR-B1* (B) and *USP18* (D) expression in testicular cell populations (cluster 1-13, 3 donors). **C)**and **E)** Analysis of *linc-UR-B1* (C) and *USP18* (E) expression in clusters of germ cells populations at different stages of prophase I (3 donors). Note that clusters 14-18 are subclusters of clusters 3 and 4 studied in(B) and (D). **F)** and **G)** Positive correlation of *linc-UR-B1* and *USP18* expression in germ cell populations (F, cluster 1-8) and in germ cells at different stages of prophase I (G, cluster 14-18). All data were retrieved from the alignment of single cell RNA-seq reads provided by (22), filtered for unique reads and expressed (± SEM) as percentage expression (normalized expression in one population *vs* all populations analyzed). Cell populations expressing *linc-UR-B1* are highlighted in red.

## DISCUSSION

USP18 determines the threshold of activation of the type I IFN signaling pathway and ultimately the amount of ISGs in a given cell. To exert its function as a negative regulator, USP18 binds to the IFN receptor/JAK complex and attenuates the magnitude of the response (2, 3, 23). The abundance of other IFN-stimulated components (*ie* STAT2/1, IRF9 and SOCS1) will also impact cell context-specific ISG activation, but in distinct and less specific manners (24). Given the critical non-redundant role of USP18, we conceived this study to assess whether USP18 can be post-transcriptionally regulated by miRNAs directed to the 3’UTR. Through the use of target prediction tools, functional validation, public data mining and correlative analyses, we identified four miRNAs (*miR-191-5p, miR-24-3p, miR-423-5p, miR-532-3p*) that directly pair to sites within the 580 nt-long 3’UTR and tune down endogenous USP18 at both mRNA and protein levels. These miRNAs are expressed at different levels in numerous immune and non-immune cell types. We do not know whether all four miRNAs collaborate to robustly target USP18. Yet, *miR-191-5p, miR-24-3p*, and *miR-532-3p* are enriched in circulating human monocytes, which we found exhibit low baseline USP18 with respect to PBL.

Monocytes are a subset of leukocytes involved in anti-microbial defenses, anti-tumor immune responses and in tissue homeostasis (25). Interestingly, Uccellini *et al*, using an ISRE-dependent reporter mouse, showed that Ly6C^hi^ inflammatory monocytes - corresponding to the classical human monocytes and representing *≈* 90% of the circulating ones - display a high basal IFN response, further enhanced upon infection (26). Under physiological conditions, circulating monocytes are primed by constitutive IFNβ, in order to be armed with adequate levels of key proteins (transcription factors, *i.e*. STATs/IRFs, some ISGs and immune response genes products) (27, 28). During acute viral infection, in some auto-immune and inflammatory diseases, monocytes encounter higher levels of IFN that in turn promotes their activation and their differentiation into dendritic-like cells with a potent antigen-presenting capacity (29). Low USP18 may be critical to maintain high responsiveness of these cells to IFN. In line with this, a recent study on *ISG15*-deficient patients showed that, among PBMCs, monocytes displayed the highest IFN signature (30). Thus, the targeting of USP18 by *miR-191-5p, miR-24-3p* and *miR-532-3p*, and possibly *miR-423-5p*, may assist the priming of monocytes towards a robust inflammatory and immune response.

The conservation analysis of the *USP18* 3’UTR sequence led to identify several duplications of the sequences spanning intron 10-exon 11. We found six copies of this sequence mapping in human chr22q11.21. We named these copies as A or B depending on the breakpoint in intron 10 (**Fig. 4B**). In chr22q11.2 there are eight LCRs, of which four (LCR22A-D) span most of chr22q11.21, a 3 Mb region that is deleted in 90% of patients affected by the 22q11.2 deletion syndrome. First described by DiGeorge in the 1960s, it represents the most frequent chromosomal microdeletion syndrome, affecting 1 per 3,000-6,000 live births. Patients show heterogeneous clinical features, multi-organ dysfunction, cognitive deficits and neuropsychiatric illness, immunodeficiency, cardiac and palatal abnormalities (19, 31). Such recurrent genomic rearrangements have fueled many studies to resolve the complex architecture of the LCR22s, identify breakpoints and study the high variability among patients (20). The LCR22A-D duplications are believed to have occurred recently, in the hominoid-lineage after their divergence from the Old World monkeys (32, 33). The highly dynamic nature of LCR22s, as of other LCRs, is believed to be a driving force in genome evolution and adaptation, as their rearrangements can give rise to new coding and non-coding genes (34, 35). Interestingly, the *USP18* protein-coding gene resides at the centromeric boundary of LCR22A. Moreover, an unprocessed pseudogene, called *USP41*, resides in the LCR22B (**Fig. 4A**). *USP41* is nearly identical to *USP18*, but lacks 5’ and 3’UTRs.

Previous in-depth studies of the LCR22s did report on several copies of *USP18* intron 10-exon 11 and suggested that *Alu* elements present at their 5’ breakpoint (**Fig. S9A**) may have driven duplication events (34, 35). Delihas referred to “*USP18*-linked” sequences as part of a repeat unit present in human and chimpanzee genomes, which includes upstream sequences related to the *GGT* (gamma-glutamyl-transferase) gene family and downstream sequences related to the *FAM230* gene family (33). Accordingly, this is the exact genomic context that surrounds the A copies of *USP18* exon 11 (**Fig. S9B**). Members of the *FAM230* family are also found downstream of the two B copies and of the *USP18* protein-coding gene, while members of the *POM121* family are found upstream (**Fig. S9B**). Altogether these observations reinforce the hypothesis that the *USP18* intron 10-exon 11 sequence was trapped, together with *FAM230* sequences, in a gene-forming element, and this event may have contributed to the generation of new genes (33).

Here, we provide the first experimental evidence that four of the six copies of the *USP18* exon 11 are embedded in expressed lincRNA genes. We refer to transcripts bearing the *USP18* exon 11, *i.e*. the entire 3’UTR, as *linc-UR-A1/3/4* and *linc-UR-B1* (**Fig. 4C, D**). By tracking the unique junction between *USP18* exon 11 and the upstream exon, we showed that *linc-UR-A* transcripts are more abundant in tissues like liver, kidney and thymus, while *linc-UR-B1* is uniquely and highly expressed in testis, notably in spermatocytes and spermatids. To our knowledge this is the first case known of a 3’UTR that is found duplicated independently from the rest of the ancestral gene and embedded in expressed non-coding RNAs. Few non-coding transcripts that bear the 3’UTR sequence of a protein-coding gene have been described. These transcripts are expressed from pseudogenes, notably *PTENP1, KRASP1* and *BRAFP1*. These latter originated by a single duplication of the ancestral protein-coding genes (coding region and UTRs), they are transcribed but not translated due to mutations creating premature stop codons or frameshifts (36, 37). Conversely, the lincRNAs that we have identified bear only the 3’UTR of *USP18*.

In recent years many non-coding RNAs have been discovered, but only few have been assigned functions (38). Moreover, a large number of lncRNA genes are transcriptionally active in testis during meiosis and spermatogenesis (39-41). The lincRNAs we describe here may represent junk DNA, proliferating in a selfish manner. Yet, evolution may have repurposed some of them to confer an advantage, be transmitted and potentially fixed in the population (42, 43). It is a fact that the inclusion of exon 11 - possibly favored by a strong constitutive acceptor site (ttctctctagGCAGGAAACT) at the intron 10-exon 11 junction - provides to these transcripts a canonical AATAAA signal for poly-adenylation. The best studied non-coding transcript bearing the 3’UTR of a protein-coding gene is *PTENP1*. This pseudogene undergoes copy number loss in human cancers, this correlating with a decrease in levels of the tumor suppressor *PTEN*. It was proposed that *PTENP1* exerts its function by sponging miRNAs, thus sustaining PTEN levels (37). Likewise, lincRNAs bearing the *USP18* 3’UTR may act as decoys titrating away *USP18*-targeting miRNAs. Interestingly, our analysis of public RNA-seq data from testicular cell subsets revealed a positive correlation between *linc-UR-B1* and *USP18* in spermatocytes and spermatids. Moreover, we found that three *USP18*-targeting miRNAs are expressed in whole testis and in germ cells. Further work is needed to investigate the possibility that in human germ cells *linc-UR-B1* and *USP18* compete for the binding of these miRNAs.

Interestingly, studies in the mouse have shown that high/continuous IFNα/β affects spermatogenesis by causing apoptosis of germ cells and induces sterility (44-46). It is therefore tempting to speculate that the maintenance of baseline *USP18* represents a recently evolved mechanism to attenuate IFN signaling and protect the precursors of spermatozoa. In this scenario human spermatocytes and spermatids may be a good shelter for viruses. Indeed, the testis is a reservoir for a number of viruses, such as Zika and Ebola viruses which can be sexually transmitted by infected men after recovery (47, 48). We recently revealed the prolonged seminal excretion of testicular germ cells persistently infected by Zika virus (Mahé *et al*, Lancet Infectious Diseases, in press). Future studies will be necessary to determine whether *linc-UR-B1* contributes to maintain *USP18* and whether this favors viral persistence in human male germ cells.

In conclusion, our work reveals the existence of miRNAs regulating USP18 through its 3’UTR, and of several lincRNAs containing the *USP18* 3’UTR. Combined, these molecules may form a non-coding network tuning USP18 levels in cell types where IFN responsiveness needs to be tightly controlled.

## MATERIALS AND METHODS

### Computational prediction of miRNAs targeting the *USP18* 3’UTR

Bibiserv (https://bibiserv.cebitec.uni-bielefeld.de/rnahybrid) and miRWalk 2.0 (http://zmf.umm.uni-heidelberg.de/apps/zmf/mirwalk2) interfaces were used. The human *USP18* 3’UTR sequence (580 nt) was downloaded from the UCSC genome browser (hg38) (https://genome.ucsc.edu/) and used as input. In Bibiserv, RNA hybrid algorithm was used with the following parameters: energy threshold = -20, no G:U in the seed, helix constraint from nt 2 to 8 (7-mer seed), approximate p-value for 3utr_human. In miRWalk, TargetScan, Pita, miRDB, RNA22 algorithms were used with the following parameters: minimum seed length 7, position 2 (nt 2) as start position of the miRNA seed, p-value 0.05.

### Monocyte isolation

PBMCs were isolated from freshly collected buffy coats obtained from healthy blood donors (Établissement Francais du Sang, Paris, France; CPSL UNT-18/EFS/041) by density gradient centrifugation using lymphocyte separation medium (Eurobio, France, Les Ulis). Monocytes were purified by positive sorting using anti-CD14-conjugated magnetic microbeads (Miltenyi Biotec, Bergisch Gladsbach, Germany). The recovered cells were >98% CD14^+^ as determined by flow cytometry with anti-CD14 Ab (Miltenyi Biotec). The CD14 negative fraction (here PBL) was also recovered for further analyses.

### Testicular cells isolation

Normal human adult testes were obtained either after orchidectomy from prostate cancer patients who had not received any hormone therapy or at autopsy. The procedure was approved by Ethics Commitee Ouest V, Rennes, France (authorization DC-2016-2783) and the French National Agency for Biomedical Research (authorization PFS09-015). The presence of full spermatogenesis was assessed by transillumination of freshly dissected seminiferous tubules. Testis fragments were frozen and stored at -80°C until RNA extraction. Pachytene spermatocytes, round spermatids, Leydig cells and peritubular cells were isolated and cultured according to previously described procedures (41) before freezing at -80°C for RNA extraction. Primary Sertoli cells were purchased from Lonza (Walkersville, MD, USA) and cultured as previously described (41).

### RNA extraction and qRT-PCR quantification

Total RNA was extracted from Trizol lysates of freshly isolated monocytes or PBL, testis explants and testicular populations, using miRNeasy mini kit (Qiagen, Germantown, MD, USA) and following the manufacturer’s instructions. Total liver RNA was purchased from Thermo Fisher scientific. Quantification and purity were assessed by a Nanodrop spectrophotometer (Nanodrop2000, Thermo Fisher Scientific). For the analysis of *USP18, IRF7, OAS1, IFIT1, STAT2, linc-UR-A1/A3/A4* and *linc-UR-B1* levels, total RNA was reverse transcribed using the high-capacity cDNA reverse Transcription kit (Applied Biosystems, Thermo Fisher Scientific). To measure the expression of *USP18, linc-UR-A1/3/4* and *linc-UR-B1* in human tissues the cDNAs from Human Immune System MTC™ Panel, Human MTC™ Panel I, Human MTC™ Panel II (Takara Bio, Mountain View, CA, USA) were used. Quantitative PCR assays were performed using the FastStart SYBR Green Master Mix (Roche) on a Step One Plus Real-Time PCR system (Applied Biosystems, Thermo Fisher Scientific). Transcript expression was normalized to the *18S* or to *ACTB* levels by using the equation 2^-ΔCt^. For miRNA expression, total RNA was reverse-transcribed using the TaqMan miRNA reverse transcription kit (Applied Biosystems, Thermo Fisher Scientific) and miRNA-specific primers (Applied Biosystems, Thermo Fisher Scientific) for hsa-*miR-24-3p*, hsa-*miR-191-5p* and hsa-*miR-532-3p*. MiRNA expression levels were then analyzed using the appropriate TaqMan miRNA assay and TaqMan Universal Master Mix II (Applied Biosystems, Thermo Fisher Scientific), according to the manufacturer’s instructions on a QuantStudio 3 Real-Time PCR system (Applied Biosystems, Thermo Fisher Scientific). The ubiquitously expressed U6b snRNA (*U6*) was quantified as above and used as endogenous control to normalize miRNA expression using the 2^-ΔCt^ formula. MiRNA qPCR-array in monocytes, testis fragments and germ cells was performed using custom 96 well plates using the Taqman Advanced technology (Applied Biosystems, Thermo Fisher Scientific). Here, *miR-425-5p* was used as endogenous control.

### Mimic reverse transfection

miRIDIAN microRNA mimics for human *miR-24-3p, miR-191-5p* and *miR-532-3p* were purchased from Horizon Discovery Ltd. A negative control - miRIDIAN microRNA Mimic Negative Control #1 - (Horizon Discovery Ltd) was used to assess the specificity of the effect driven by the specific miRNA sequences. Mimic reverse transfection (50 nM) was performed in HeLa S3 cells cultured in DMEM supplemented with 10% heat inactivated FCS and antibiotics. Transfection was carried out by using Lipofectamine RNAiMAX (Thermo Fisher Scientific) according to the manufacturer’s instructions. Briefly, the mimic-RNAiMAX complexes were prepared in Opti-MEM serum-free medium by mixing 50 nM of mimic with a dilution of 1:1000 of lipofectamine RNAiMAX (calculated on the final volume of the culture). After 15 min at room temperature, the complexes were distributed in a p24-(for RNA isolation and luciferase assays) or a p60-(for protein extraction) well plate and 1×10^5^ or 9×10^5^ cells respectively were plated on top of the complex. After 48h of transfection, cells were harvested and lysed for RNA isolation, luciferase assay or protein extraction.

### Identification and analysis of duplicated *USP18* sequences

To identify *USP18* copies, we performed BLAT (https://genome.ucsc.edu/cgi-bin/hgBlat) analysis using human *USP18* gene sequence. The sequence was downloaded from UCSC genome browser (hg38) (https://genome.ucsc.edu/). We selected only sequences with more than 95 % identity to *USP18* sequence and longer than 100 bp. To find whether the *USP18* copies overlapped genes, we looked at the coordinates of these repeats on the UCSC genome browser (hg38). The genes overlapping *USP18* copies were identified, and their genomic sequences and predicted mRNA sequences were downloaded. To measure the expression of *USP18* related (*UR*) genes, we designed specific primers, targeting the unique exon-exon11 junctions to avoid detection of *USP18* mRNA. Next, we performed qPCR analysis as described above. All qPCR products were verified by sequencing.

### Analysis of public available datasets

Data on miRNA expression were obtained from the FANTOM5 dataset (https://fantom.gsc.riken.jp/5/suppl/De_Rie_et_al_2017/vis_viewer/#/human) and from (18) (Cohort Roche, GEO accession: GSE28492). The FANTOM5 dataset was downloaded and analyzed using Qlucore Omics Explorer software. RNA-seq datasets covering a panel of 32 tissues (https://www.ebi.ac.uk/gxa/experiments/E-MTAB-2836/Results) were used in **Fig. S3B**. PolyA+ RNA-seq track from testis provided by the ENCODE project (GEO accession: GSM2453457-GSM2453458) and shown in **Fig. S4** were visualized using autoscale mode in IGV browser. The data of the single cell RNA-seq on testis used in **Fig. 6** were retrieved from (22). Reads were aligned on the hg38 assembly of the human genome and only unique reads were filtered.

### Luciferase reporter assay

For cloning the *USP18* 3’UTR in the psiCHECK2 vector (Promega, Madison, WI, USA), the first 559nt of *USP18* 3’UTR were amplified from cDNA of HeLa S3 treated with IFN using the primers USP183UTR_XhoI_FW and USP183UTR_NotI_RV. Subsequently, the amplicon and psiCHECK2 vector were digested by Xho1 and Not1 and ligated. The resulting plasmid (psiCHECK2-*USP18* 3’UTR) was mutated (nt 2 to 6 of the seed-matched sequence) in the 3’UTR binding sites of *miR-24-3p, miR-191-5p* and *miR-532-3p*. Mutations were generated in the psiCHECK2-*USP18* 3’UTR plasmid using QuikChange XL site-directed mutagenesis kit (Aligent Technologies, Santa Clara, CA, USA) and specific primers for each site (24-3p_BSmut_FW/RV, 191-5p_BSmut_FW/RV, 532-3p_BSmut_FW/RV). All new plasmids were sequenced and named as BSmut *miR-24-3p, miR-191-5p* and *miR-532-3p*. For the luciferase reporter assay, 1×10^5^ HeLa S3 cells/well were reverse transfected in a 24-well plate (see section on *Mimic reverse transfection*). After 24h, plasmid transfection was carried out using FuGENE HD (Promega) according to the manufacturer’s instructions. Three biological replicates were prepared for each condition. Cells were lysed 24 hr after transfection and analyzed with Dual-Luciferase® Reporter Assay System (Promega) according to the manufacturer’s instructions. Firefly luciferase was measured as normalizer.

### Western blot analysis and antibodies

Cells were lysed in modified RIPA buffer (50 mM Tris-HCl pH 8, 200 mM NaCl, 1% NP40, 0.5% DOC, 0.05% SDS, 2mM EDTA) with 100 mM PMSF and phosSTOP and a cocktail of antiproteases (Roche). A total of 40 μg proteins was separated by SDS-PAGE and analyzed by western blot. Membranes were cut horizontally according to molecular size markers, and stripes were incubated with different Abs. Immunoblots were analyzed with the ECL Western blotting Reagent (Pierce) or the more sensitive Western Lightning Chemiluminescence Reagent Plus (PerkinElmer) and bands were quantified with Fuji LAS-4000. For reprobing, blots were stripped in 0.2 M glycine (pH 2.5) for 20 min at rt. The following Abs were used: rabbit anti-USP18 (D4E7, Cell Signaling Technology, Beverly, MA) and mouse anti-actin B (Sigma-Aldrich, St. Louis, MO, USA).

### 3’ of cDNA ends (3’RACE) and PCR

3’ RACE analysis was performed on testis and liver RNA using the 3’ RACE System kit (Thermo Fisher Scientific) according to manufacturer’s instructions. PCR was performed with Platinum SuperFi II DNA Polymerase (Thermo Fisher Scientific).

### Primers

For the sequences of all primers used refer to **Table S2**.

## Supporting information

Rubino et al_Supplementary information

## ACKNOWLEDGMENTS

We wish to thank P. Miesen for help in initial prediction analyses of miRNAs targeting USP18 3’UTR; J. Guo for assisting us in retrieving scRNA-seq dataset on the Human Testis Cell Atlas; the Flow Cytometry Platform of Institut Pasteur and S. Meunier for technical assistance for cytometric analyses; the members of the Unit of Cytokine Signaling, M. Livingstone, G. Uzé, V. Libri and N. Jouvenet for advice, discussions and support; M. C. Gauzzi for critical reviewing of the manuscript.

## AUTHOR CONTRIBUTIONS

ER and MC designed and performed experiments; ÖV, LV and GG performed experiments under supervision; ER and MC analyzed the data, NT and ER performed and analyzed the RNA-seq and sequence conservation data; AL, AR and NDR provided testis samples and performed experiments on purified testicular populations; NDR and FM provided expert advice on experiments; ER, MC and SP wrote the manuscript; all authors revised the manuscript; SP supervised the work.

## FUNDING

Research in the Unit of Cytokine Signaling is funded by the Institut Pasteur, the Fondation pour la Recherche Médicale (Equipe FRM DEQ20170336741) and the Institut National de la Santé et de la Recherche Médicale (INSERM). ER was supported by Sorbonne Université and by FRM. MC was supported by the FRM grant above. GG and ÖV were supported by the Erasmus plus programme of the EU commission.

## Notes

### Competing Interest Statement

The authors have declared no competing interest.

