## Supplementary material for "Human USP18 is regulated by miRNAs *via* the 3’UTR, a sequence duplicated in lincRNA genes residing in chr22q11.21": Rubino et al_Supplementary information

**This PDF file includes:**

Figures S1 to S9
Tables S1 and S2

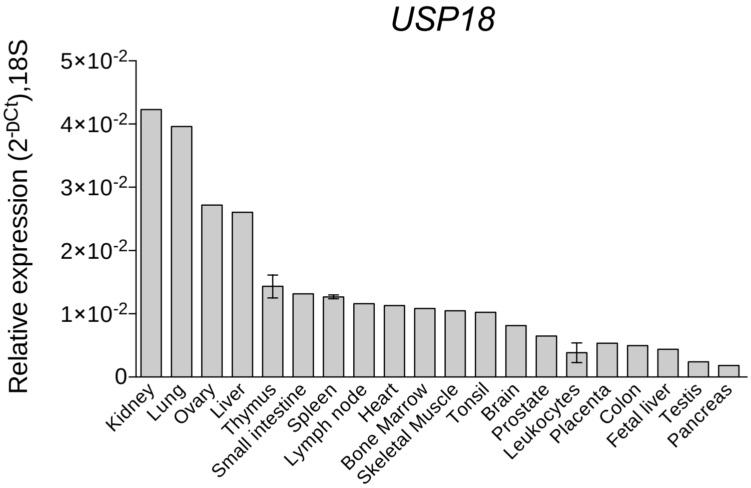


**S1 Figure. *USP18* is widely expressed in human tissues.**

Expression of *USP18* in 20 human tissues (Human Immune System MTC™ Panel, Human MTC™ Panel I, Human MTC™ Panel II). Results shown as expression ($2^{-\Delta Ct}$) relative to *ACTB*, used as housekeeping gene. SEM is shown for tissues that are present in Human Immune System MTC™ Panel and Human MTC™ Panel I.

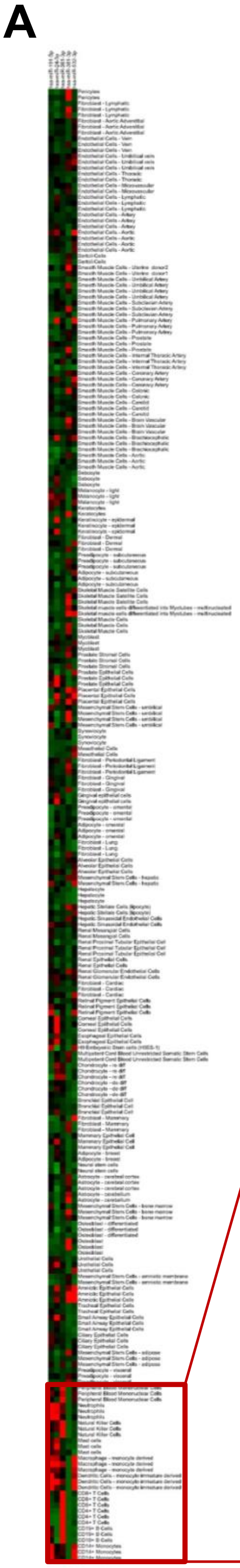


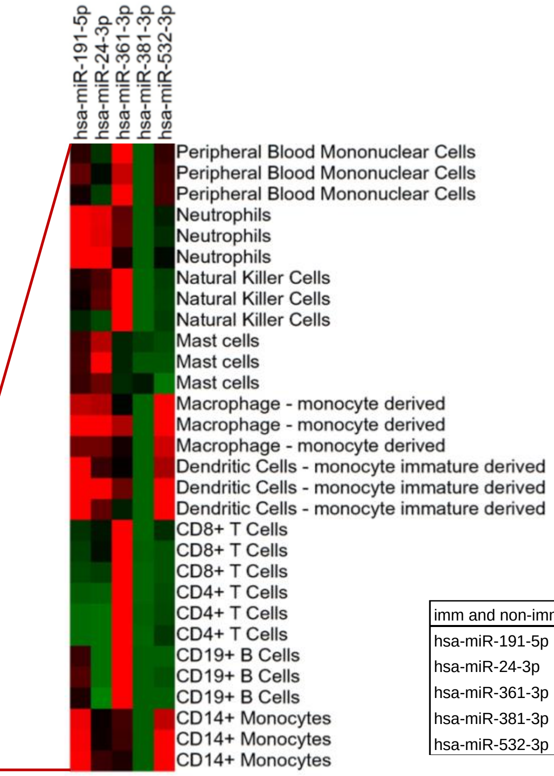

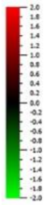


| imm and non-imm | p-value | q-value |
|---|---|---|
| hsa-miR-191-5p | 1.26E-18 | 4.40E-18 |
| hsa-miR-24-3p | 0.000611121 | 0.001069462 |
| hsa-miR-361-3p | 5.19E-38 | 3.64E-37 |
| hsa-miR-381-3p | 9.86E-06 | 2.30E-05 |
| hsa-miR-532-3p | 0.01453944 | 0.020355216 |

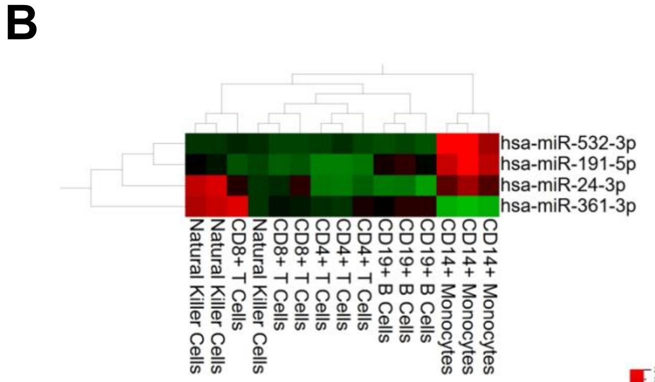


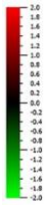

| circulating PBMCs | p-value | q-value |
|---|---|---|
| hsa-miR-191-5p | 2.55E-07 | 1.02E-06 |
| hsa-miR-24-3p | 0.003559705 | 0.004746273 |
| hsa-miR-361-3p | 0.025307131 | 0.025307131 |
| hsa-miR-532-3p | 8.89E-07 | 1.78E-06 |

**S2 Figure. *miR-191-5p*, *miR-24-3p* and *miR-532-3p* are enriched in immune cells and particularly in monocytes.**

A) Heat map visualization of the data in Fig. 2A. Differential expression of *miR-191-5p*, *miR-24-3p*, *miR-361-3p*, *miR-381-3p* and *miR-532-3p* in 90 human cell types (immune and non-immune) is shown. Immune cells are delimited by a red box and zoomed. p-values and q-values (ANOVA FDR adjusted p-value) are shown on the right of the heat map for all significant miRNAs.

B) Heat map visualization of the data in Fig. 2B. Differential expression of *miR-191-5p*, *miR-24-3p*, *miR-361-3p* and *miR-532-3p* in circulating PBMCs is shown. p-values and q-values (ANOVA FDR adjusted p-value) are shown below the heat map for all significant miRNAs.

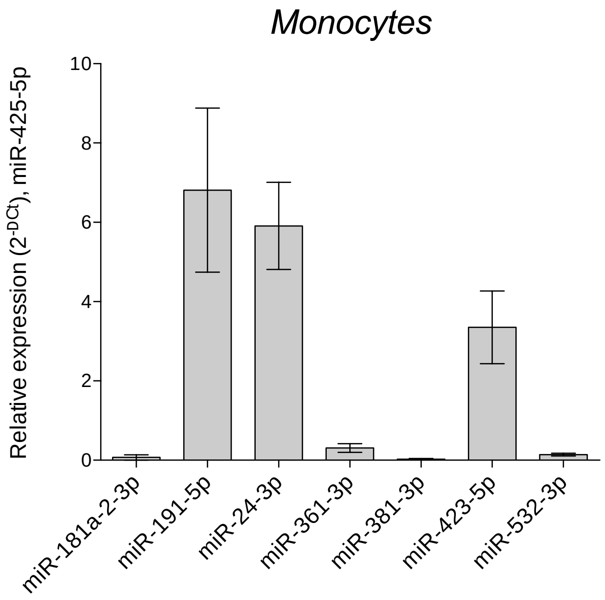


**S3 Figure. *USP18*-targeting miRNAs in monocytes.**

The expression of the indicated *USP18*-targeting miRNAs in monocytes was measured by miRNA qPCR-array. *miR-425-5p* was used as normalizer

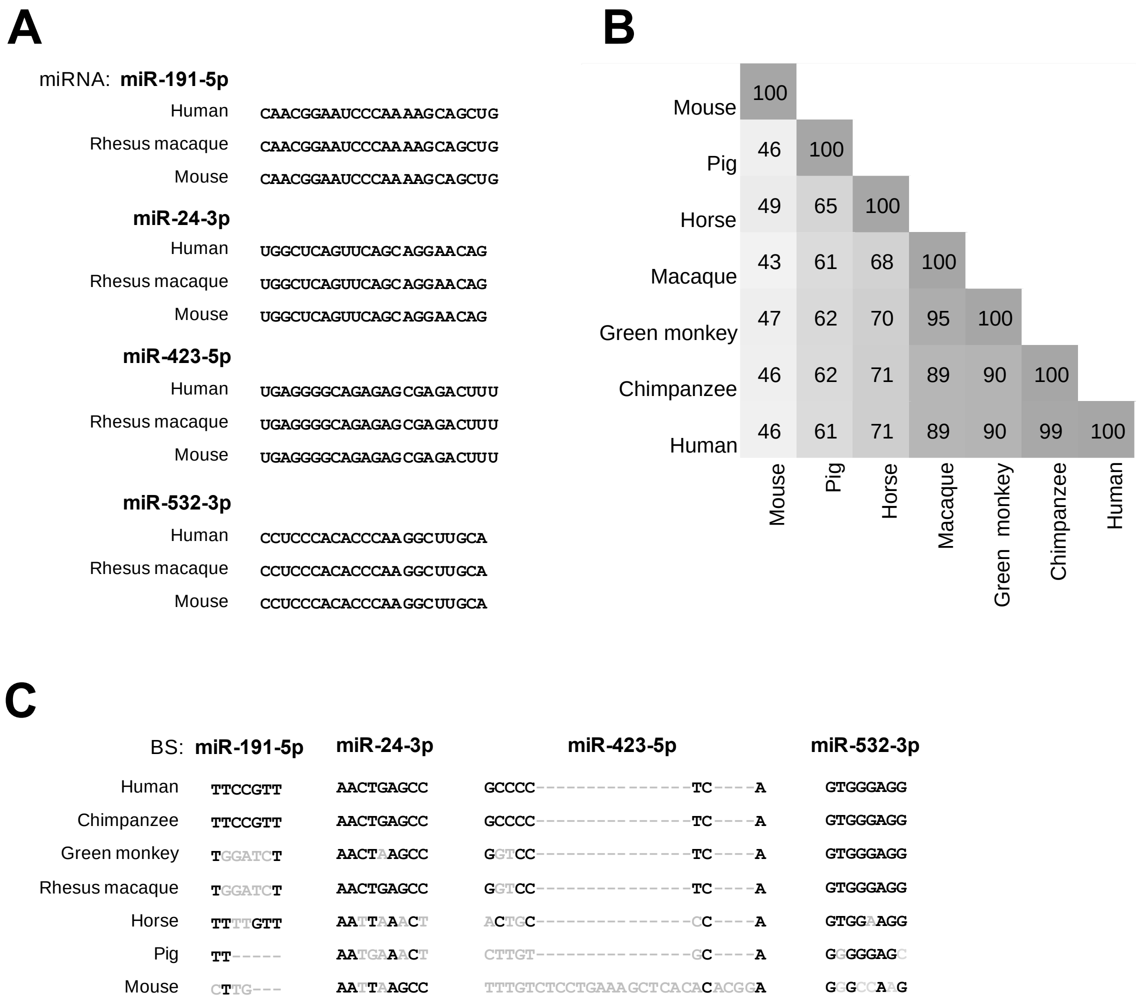


**S4 Figure. Conservation of *USP18* 3’UTR and miRNA binding sites.**

A) Conservation of *miR-191-5p*, *miR-24-3p*, *miR-423-5p* and *miR-532-3p* mature sequence in rhesus macaque (non-human primate) and mouse. Conserved nucleotides in black. Sequences retrieved from mirbase (www.mirbase.org).

B) Identity matrix of *USP18* 3’UTR in mammals. Each number indicates the percentage of identity between two species. Shades of gray indicate level of conservation (low in light gray; high in dark gray). *USP18* 3’UTR sequences were downloaded from UCSC (https://genome.ucsc.edu). Sequences were aligned using Clustal Omega (https://www.ebi.ac.uk/Tools/msa/clustalo).

C) Conservation of the seed-matched region of the binding site (BS) of *miR-191-5p*, *miR-24-3p*, *miR-423-5p* and *miR-532-3p* on the *USP18* 3’UTRs of mammals. Conserved nucleotides in black; non-conserved nucleotides in gray.

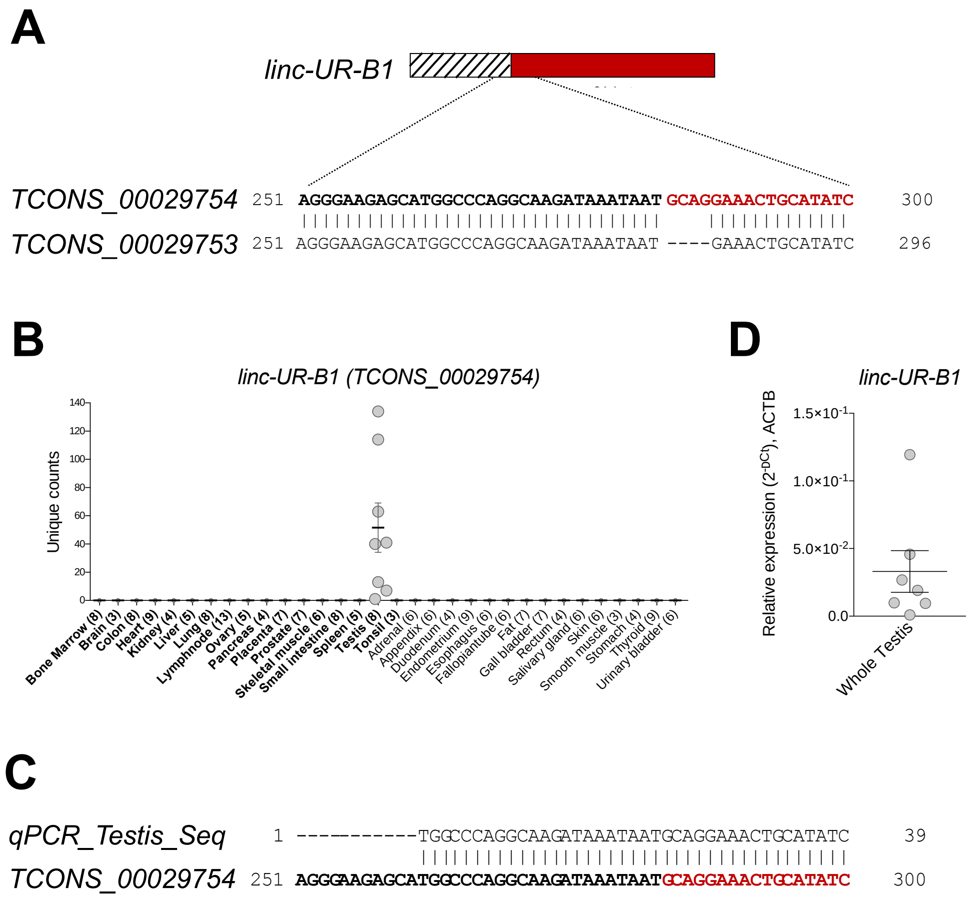


**S5 Figure. *TCONS_00029754* is the *linc-UR-B1* isoform expressed in testis.**

A) Alignment of the exon-exon junction of *TCONS_00029754* and *TCONS_00029753* (putative *linc-UR-B1* isoforms). In the alignment, gaps are indicated with dashes. Note that an ATG codon in frame with the last 46 coding nucleotides of *USP18* is present only in *TCONS_00029754*.

B) Expression of *TCONS_00029754*, here called *linc-UR-B1*, in 32 human tissues (RNA-seq data from https://www.ebi.ac.uk/gxa/experiments/E-MTAB-2836/Results), shown as unique counts. In bold the tissues in common with tissues in the panel analyzed by qPCR in Fig. 4G. The other tissues are unique to this dataset. The number of donors is shown in brackets. *linc-UR-B1* detected only in testis (8 donors).

C) Sequencing of the qPCR product obtained from testis cDNA in Fig. 4G, revealed that the isoform of *linc-UR-B1* expressed in testis is *TCONS_00029754*.

D) Expression of *linc-UR-B1* in testis fragments (7 donors), measured by qPCR. Expression is shown as relative ($2^{-\Delta Ct}$) to *ACTB*. The donor with the lowest expression of *linc-UR-B1* had impared spermatogenesis, while the other six donors

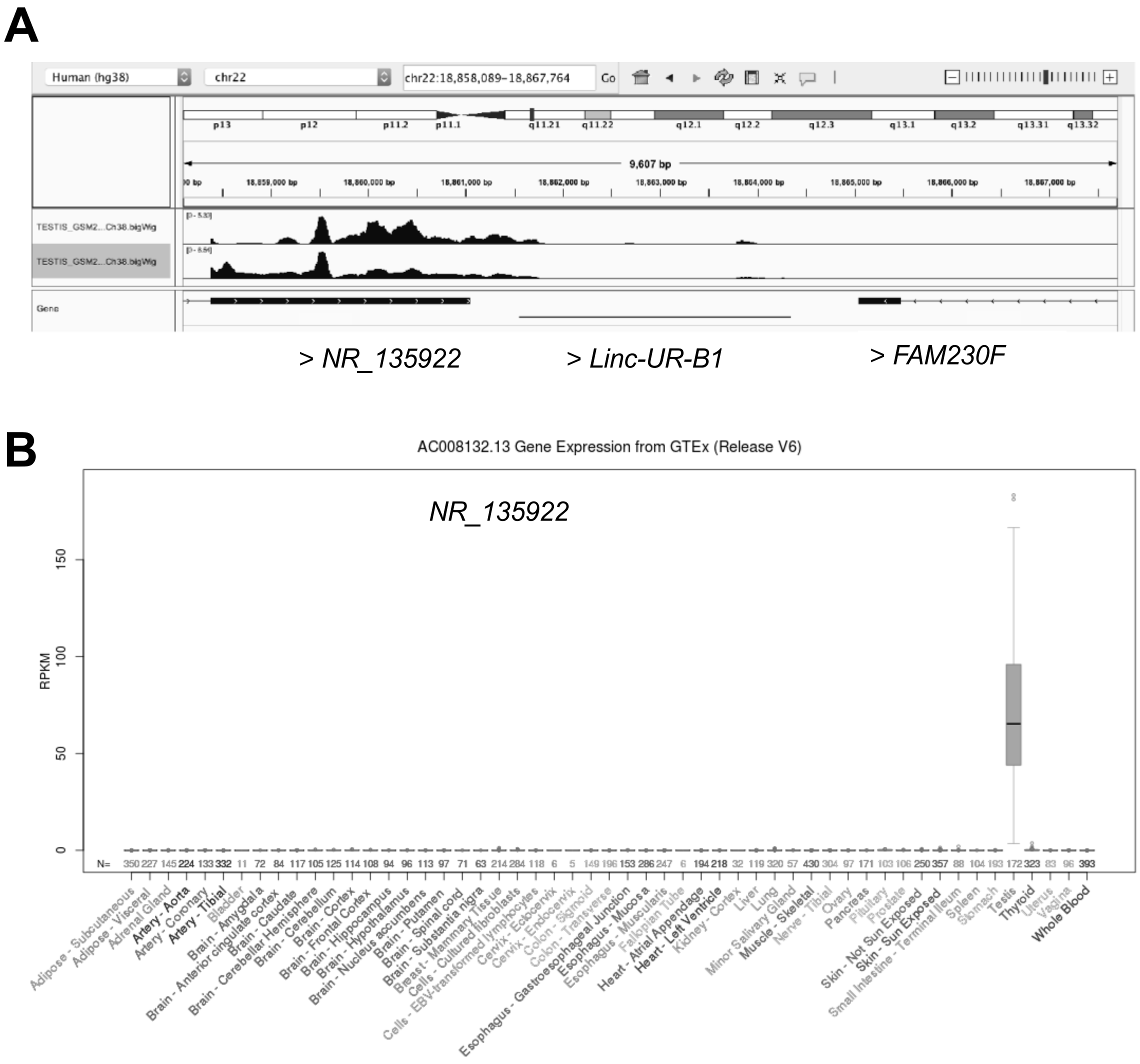


**S6 Figure. The *NR_135922* pseudogene is uniquely expressed in testis.**

A) PolyA+ RNA-seq (+ strand) track from testis provided by the ENCODE project (GSM2453457 and GSM2453458) were visualized using auto-scale mode in IGV browser (hg38 genome assembly). Genes are shown with their genomic orientation ( > for + strand, < for – strand). *NR_135922* and the annotated intergenic region upstream of *linc-UR-B1* is covered by reads.

B) Expression of *NR_135922* in GTEx data, retrieved from UCSC. *NR_135922* gene is under the name AC008132.12 and is expressed only in testis.

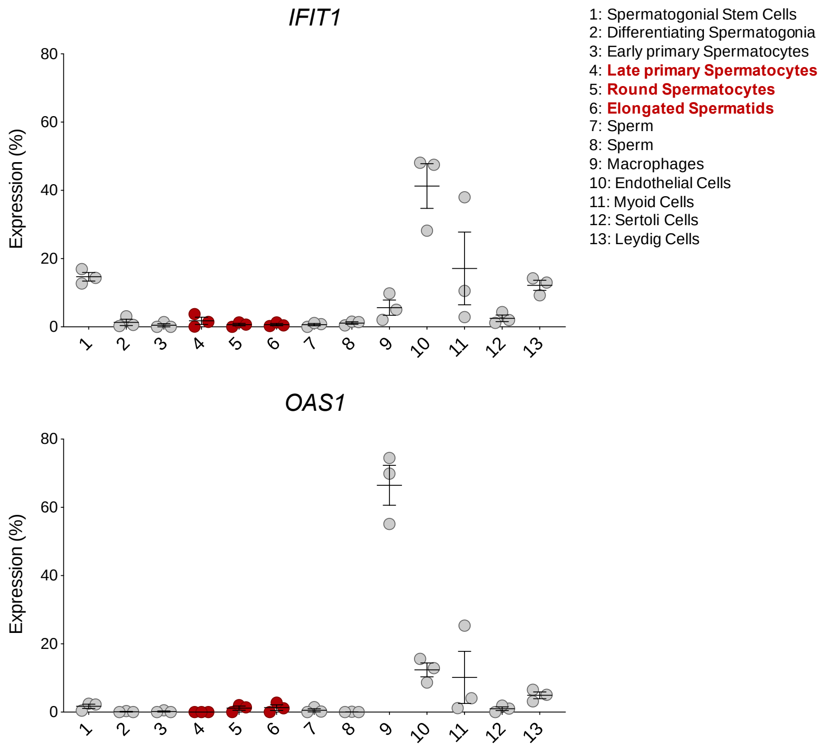


**S7 Figure. No or low expression of ISGs in germ cells.**

Analysis of *IFIT1* and *OAS1* in testicular cell populations (cluster 1-13, 3 donors). All data were retrieved from the alignment of single cell RNA-seq reads provided by [22], filtered for unique reads and expressed as percentage expression (normalized expression in one population *vs* all populations analyzed). Cell populations expressing *linc-UR-B1* are highlighted in red.

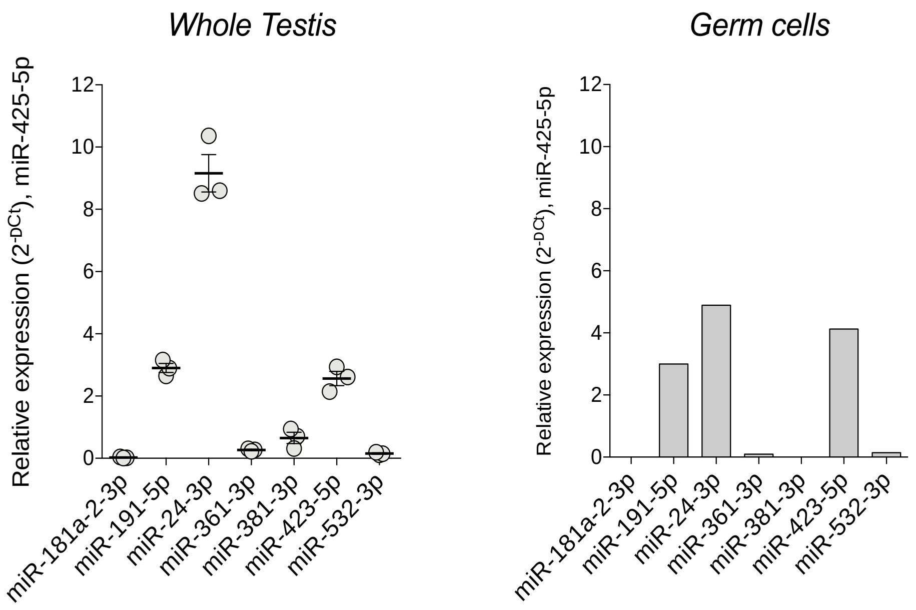


**S8 Figure. *USP18*-targeting miRNA are expressed in whole testis and germ cells.** The expression of *USP18*-targeting miRNAs in testis fragments (left panel) and on purified germ cells (right panel) was measured by miRNA qPCR-array. *miR-425-5p* was used as normalizer.

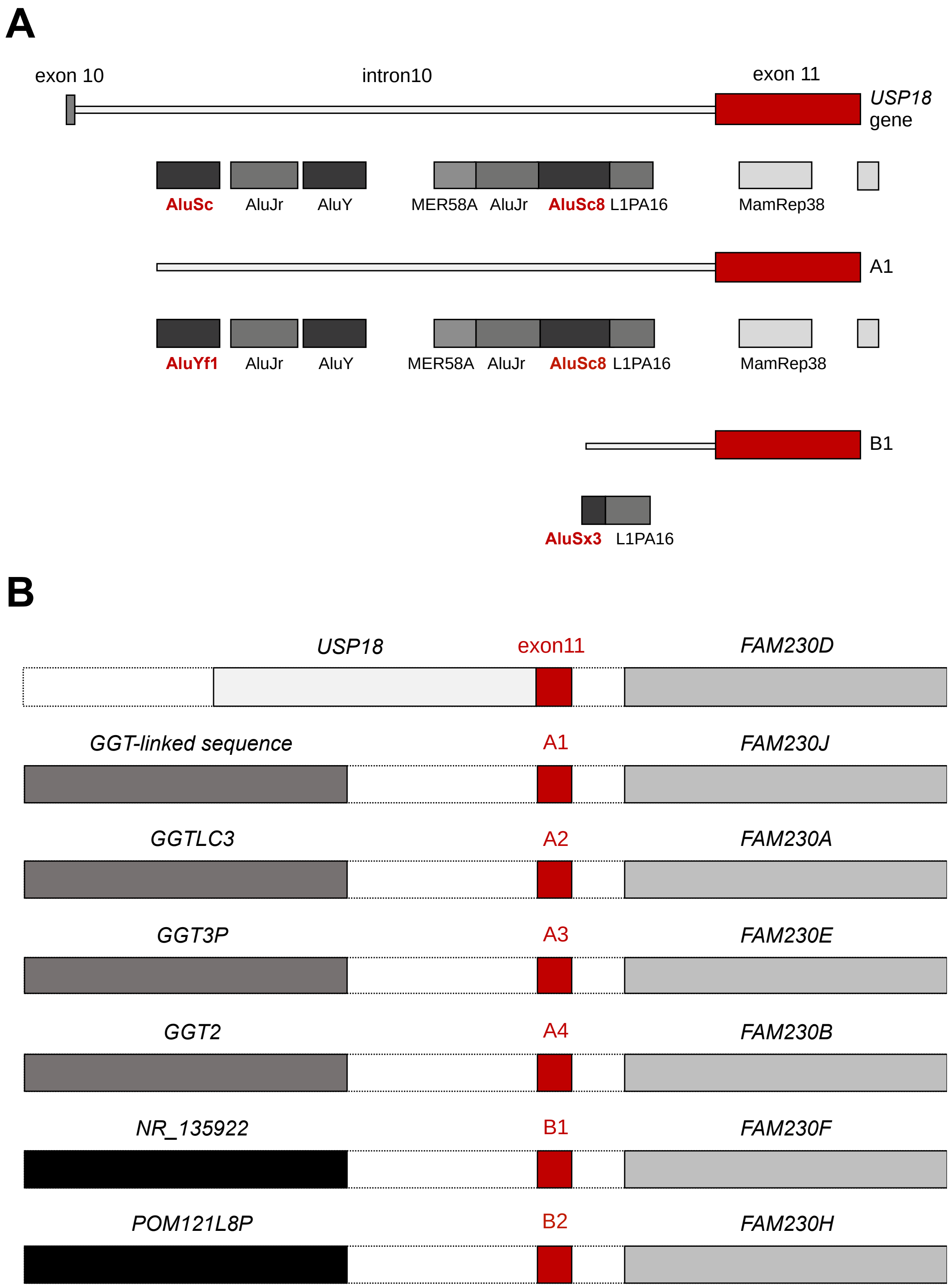
A
exon 10
intron10
exon 11
USP18 gene
AluSc
AluJr
AluY
MER58A
AluJr
AluSc8
L1PA16
MamRep38
A1
AluYf1
AluJr
AluY
MER58A
AluJr
AluSc8
L1PA16
MamRep38
B1
AluSx3
L1PA16
B
USP18
exon11
FAM230D
GGT-linked sequence
A1
FAM230J
GGTLC3
A2
FAM230A
GGT3P
A3
FAM230E
GGT2
A4
FAM230B
NR_135922
B1
FAM230F
POM121L8P
B2
FAM230H

**S9 Figure. *USP18* intron 10-exon 11 copies contain *Alu* elements at their breakpoints and are part of repeated gene blocks.**

A) Repeated elements present in the A and B copies (A1 and B1 are shown as representatives). The *Alu* sequences at the breakpoint of the A and B copies are shown in red. Exons are shown as boxes, intron as thick lines. Sequences identical to *USP18* exon 11 are in red.
B) *USP18* exon 11, A copies and B copies are shown in red. *FAM230*-linked sequences/genes are found downstream of A and B copies. GGT-linked sequences/genes are shown upstream of A copies. POM121-related sequences/genes (*POM121L8P* and *NR_135922*) are found upstream of the B copies. Genes/sequences are shown with their genomic orientation ( > for + strand, < for – strand).

**Table S1. Copies of the sequences annotated as exon 1 of *linc-UR-A1/3/4* and *linc-UR-B1.***

| Coordinates [a] | Strand [b] | Identity % (score) [c] |
|---|---|---|
| ***linc-UR-A*** | | |
| **chr22:18769207-18769785** [d] | - | 100 (578/578) |
| **chr22:18349936-18350514** | + | 100 (578/578) |
| **chr22:21203454-21204032** | - | 99.9 (576/578) |
| **chr22:18511886-18512464** | + | 99.9 (576/578) |
| chr20:23980580-23981153 | - | 96.2 (536/578) |
| chr13:18248901-18249486 | - | 96.6 (533/578) |
| chr22:22651789-22652345 | + | 96.6 (531/578) |
| chr22:24632871-24633427 | + | 96.6 (525/578) |
| chr22:24244569-24245122 | - | 96.6 (490/578) |
| ***linc-UR-B1*** | | |
| **chr22:18861451-18861733** | + | 100 (283/283) |
| chr22:21297396-21297678 | + | 100 (283/283) |
| chr22:21112840-21113122 | - | 99.7 (281/283) |
| chr6:162068502-162068656 | - | 82.6 (101/283) |

[a] Blat was performed on human genome using *linc-UR-A1* or *linc-UR-B1* annotated exon1 sequence as input.
[b] Refers to the genomic DNA strand
[c] Hits with identity > 80% and length > 100bp are shown.

**Table S2. Sequences of the primers used.**

| Primers | Sequences 5’ – 3’ |
|---|---|
| USP18_qPCR_FW | ACTCCTTGATTTGCGTTGAC |
| USP18_qPCR_RV | TTTCCCACGGGTCTTCTT |
| OAS1_qPCR_FW | TTGACTGGCGGCTATAAACC |
| OAS1_qPCR_RV | TGGGCTGTGTTGAAATGTGT |
| IRF7_qPCR_FW | GGGTGTGTCTTCCCTGGATA |
| IRF7_qPCR_RV | GCTCCATAAGGAAGCACTCG |
| IFIT1_qPCR_FW | TCTCAGAGGAGCCTGGCTAA |
| IFIT1_qPCR_RV | TCAGGCATTTCATCGTCATC |
| STAT2_qPCR_FW | TATCACAGCCAGTGCCAGAG |
| STAT2_qPCR_RV | CTGATTCCCATCCTTGGAGA |
| 18S_qPCR_FW | CATGGCCGTTCTTAGTTGGT |
| 18S_qPCR_RV | CGCTGAGCCAGTCAGTGTAG |
| ACTB_qPCR_FW | TACAGCTTCACCACCACGG |
| ACTB_qPCR_RV | TGCTCGAAGTCCAGGGCGA |
| USP183UTR_XhoI_FW | CGCGCTCGAGTGGAAATGCCCAAAACCTTC |
| USP183UTR_NotI_RV | GCGCGCGGCCGCTCATGACTGTGTTTATCAC |
| 532-3p_BSmut_FW | CCAGTGGGGAGAGCAGTGGCAGTCCCTCGCATCTGGGGGC |
| 532-3p_BSmut_RV | GCCCCCCAGATGCGAGGGACTGCCACTGCTCTCCCCACTGG |
| 24-3p_BSmut_FW | GTTACATATTTTGATAATATCCCTAATTATAAATAAGCGAGTGTTATATAGTTTGAAAACAATGCTTCTCCTCATTGCA |
| 24-3p_BSmut_RV | TGCAATGAGGAGAAGCATTGTTTTCAAACTATATAACACTCGCTTATTTATAATTAGGGATATTATCAAAATATGTAAC |
| 191-5p_BSmut_FW | AAGACTCCGTAGATCCAGGATGCCTAATGGAAAATGACAGCGTGTCAATCTCTG |
| 191-5p_BSmut_RV | CAGAGATTGACACGCTGTCATTTTCCATTAGGCATCCTGGATCTACGGAGTCTT |
| 3UTR_GSP1 | AGTTGTATAATACTGAAG |
| A_FW1 | CTGTTGCTGCTGACTCCAAG |
| A_RV1 | TCCGTAGATCCAGGAACGGAA |
| A_FW2 | TGAGGCATGAGTTTGGCCAC |
| B_FW1 | TGGCCCAGGCAAGATAAATA |
| B_RV1 | TGGTGAAAGCATCCATTCTG |
| B_FW2 | CATTGATTACGACTTCCCTTCACCAC |
| B_FW3 | CTTCTAACCCAGAGAACACAGC |
| B_FW4 | CCTGGTGTCCATGCTTTTGTGA |
| B_FW5 | CTCAGTTTCTGGTTACATCTGA |
| B_FW6 | GCTGGCACTGCGAGCAATATA |
| 3UTR_RV | TGAGGGGCCTCATGGTTACA |
